# An effector deletion leads to the breakdown of partial grapevine resistance to downy mildew

**DOI:** 10.1101/2023.08.17.553663

**Authors:** Manon Paineau, Andrea Minio, Pere Mestre, Frédéric Fabre, Isabelle D. Mazet, Carole Couture, Fabrice Legeai, Thomas Dumartinet, Dario Cantu, François Delmotte

## Abstract

Grapevine downy mildew, caused by the oomycete *Plasmopara viticola*, is a globally destructive disease that particularly affect the Eurasian wine grape *V. vinifera*. While genetically resistant varieties are becoming more accessible, populations of *P. viticola* are demonstrating rapid adaptability, successfully overcoming these resistances. Here we aimed to identify the avirulence genes involved in the interaction with the Rpv3.1-mediated resistance in grapevine. We sequenced the full genome of 136 *P. viticola* strains sampled in a natural population of Bordeaux (France) and characterized their development on both resistant and sensitive cultivars. The genome-wide association study allowed the identification of a genomic region associated with the breakdown of Rpv3.1 grapevine resistance (avrRpv3.1 locus). A diploid-aware reassembly of the *P. viticola* INRA-Pv221 genome allowed to detect structural variations in this locus, including a major 30 Kbp deletion. At the avrRpv3.1 locus, virulent *P. viticola* strains presented deletion on both haplotypes indicating that avirulence is recessive. The deletion involves two closely-related genes that encode proteins containing 800-900 amino acids with a signal peptide. The structure of the predicted proteins contains repeats of the LWY-fold structural modules, typical of oomycete effectors. Moreover, when these proteins were transiently expressed, they induced cell death in grapevines carrying Rpv3.1 resistance, confirming their avirulence nature. The first description of candidate effectors of *P. viticola* involved in the interaction with resistance gene provides valuable insights into the genetic mechanisms that enable *P. viticola* to adapt to grapevine resistance, laying a foundation for developing strategies to manage this damaging crop pathogen.

## Introduction

Grapevine downy mildew, caused by the obligate biotrophic oomycete *Plasmopara viticola* (Berk. & M. A. Curt.) Berl. & De Toni, is one of the most destructive diseases world-wide (1). The disease was endemic in North America (2, 3), where native grape species had developed genetic resistance, and it was accidentally introduced to southwest France in the 1870s from where it rapidly spread throughout Europe (4, 5). The Eurasian wine grape *Vitis vinifera* L. is highly susceptible to this disease and disease control is mostly achieved through the use of fungicides. Grapevine breeding programs around the world have been producing new cultivars genetically resistant to the disease by introgressing disease resistance *loci* from wild grape species. Resistance to downy mildew found in grapevine is mainly partial and caused by Resistance-genes (R-gene) that, depending on the resistance source, may explain up to 80% of downy mildew infection on the plant (6, 7). Over 30 downy mildew resistance *loci* have been identified in American and Asian *Vitis* species (8), but only few of them are currently being utilized in breeding programs : Rpv1 (9), Rpv3 (10–12), Rpv10 (13), and Rpv12 (6). Many of these cultivars are now commercially available and gaining popularity among growers, especially following the promotion of agroecological transition and pesticide use restrictions imposed by the European Union (14).

*Plasmopara viticola* possesses a remarkable ability to evolve, as demonstrated by its rapid adaptation to synthetic fungicides (15–17). In addition to large population size and an obligatory sexual cycle (18), *P. viticola* boasts a highly repetitive genome, altogether granting it a great evolutionary potential (19, 20). Consistent with this and despite a limited deployment of disease-resistant grape cultivars in vineyards, breakdown of resistance has been reported, and some *P. viticola* strains are already able to simultaneously overcome several resistance loci, including Rpv3.1, Rpv3.2, Rpv10, and Rpv12 (17, 21–26). The ability of *P. viticola* populations to rapidly adapt to resistant grapevines is concerning because of the substantial breeding efforts required to identify resistance sources and develop new resistant cultivars. Minimizing the risk of grapevine resistance genes depletion requires a better understanding on the genetic mechanisms responsible for pathogen adaptation to host resistance.

Grapevine resistance to downy mildew is marked by the triggering of hypersensitivity responses (HR) upon infection by *P. viticola*, suggesting a gene-for-gene interaction between the plant and the pathogen. Most genetic factors of grape conferring resistance to downy mildew identified until now are located in genomic regions rich in Nucleotide Binding Domain and Leucine-Rich Repeat (NBS-LRR) genes (7, 27, 28). The presence of NBS-LRR genes was confirmed by the cloning of the loci causing Rpv1 and Rpv3 resistance (29, 30). NBS-LRR proteins are involved in the recognition of specialized pathogen effectors. In oomycetes, the RXLR family is the largest and most studied family of cytoplasmic effectors. RXLR effectors are characterized by the presence of a signal peptide, a RXLR-EER motif at their N-terminal sequence and one or more WY-domains, a common fold found only in this family of proteins (31). Structural studies revealed an additional fold present in RXLR effectors, the LWY domain, which is often present in tandem repeats, conferring structural and functional modularity (32). Related candidate effector proteins that possess a signal peptide and carry WY-domains and the EER motif, but lack an RXLR motif, have been described in several downy mildews species (33–36). To date, all the effectors of *P. viticola* that have been investigated were discovered through *in silico* predictions (19, 37–42). While these studies have provided valuable insights into the mechanisms employed by the pathogen to infect its host, none of them have been reported to be directly related to a gene-for-gene interaction with resistance loci. Therefore, the specific effectors targeted by grape resistance genes remain entirely unknown to date.

Rpv3.1 is the most exploited resistance in viticulture (12). This resistance has been introduced from an American grape species into the crop germplasm at the end of the 19th century. It is present in most of the French-American hybrids and in many modern varieties resistant to downy mildew. In response to *P. viticola* infection, Rpv3.1 triggers effector-triggered immunity and localized necrosis in grapevine leaves (11). The causal factor for this resistance was recently mapped to a single locus of grapevine genome containing two TIR-NB-LRR (TNL) genes that originated from a tandem duplication (interleukin-1 receptor-nucleotide binding-leucine-rich repeats) (30). On the pathogen side, rapid adaptation to Rpv3.1-mediated resistance has been reported in several geographically distant populations of *P. viticola* over the past decades (17, 21–24, 26, 43). Overall, these findings suggests a gene-for-gene interaction between TNL genes of the plant and an undescribed avirulence gene of the pathogen.

In recent years, genome-wide association studies (GWAS) have become increasingly popular for identifying virulence/avirulence factors in fungal plant pathogens (44–51). In oomycetes, while the method has shown success in detecting the genomic regions underlying fungicide resistance and mating-type (20, 52), it has not yet been employed for discovering avirulence genes. In this study, we used GWAS to identify the avirulence genes interacting with Rpv3.1-mediated resistance of grapevine. To this end, we sampled 136 strains of *P. viticola* from a natural population in Bordeaux (France) and characterized their development on both resistant (Rpv3.1) and susceptible grapevines. By sequencing the genome of these strains, we were able to identify candidate effectors and the genomic event responsible for the breakdown of Rpv3.1-mediated resistance.

## Results

### Localization of the virulence locus by GWAS

Sequence data was obtained for 123 strains (over the 136 sequenced). A total of 1,851,765 SNPs were retained, resulting in an average density of 20.8 SNPs / kbp on the 359 scaffolds of the *P. viticola* reference genome. To assess the phenotypic variability of the 123 *P. viticola* strains, we evaluated the percentage of leaf disc area covered by sporulation in a cross-inoculation experiment between the 123 *P. viticola* strains and two grapevine varieties: Cabernet Sauvignon (susceptible, Rpv3.1-), and Regent, carrying the resistance locus Rpv3.1 (Rpv3.1+) (Figure 1A and B). On Cabernet Sauvignon, the strains displayed an average of sporulation of 10.4% (sd = 3.92) (in gray on Figure 1B). On Regent, the average sporulation area of the strains was 6.91% (sd=6.41) (in orange on Figure 1B). While the distribution of sporulation area of the strains on Cabernet Sauvignon is centered around the mean, the sporulation variability among strains observed on Regent ranged from 0.01% to 29.33% and follows a bi-modal distribution (Figure 1A): a first group of strains displaying around 2.5% of sporulation area on Regent and a second group displaying around 14% of sporulation area.

**Fig. 1.**
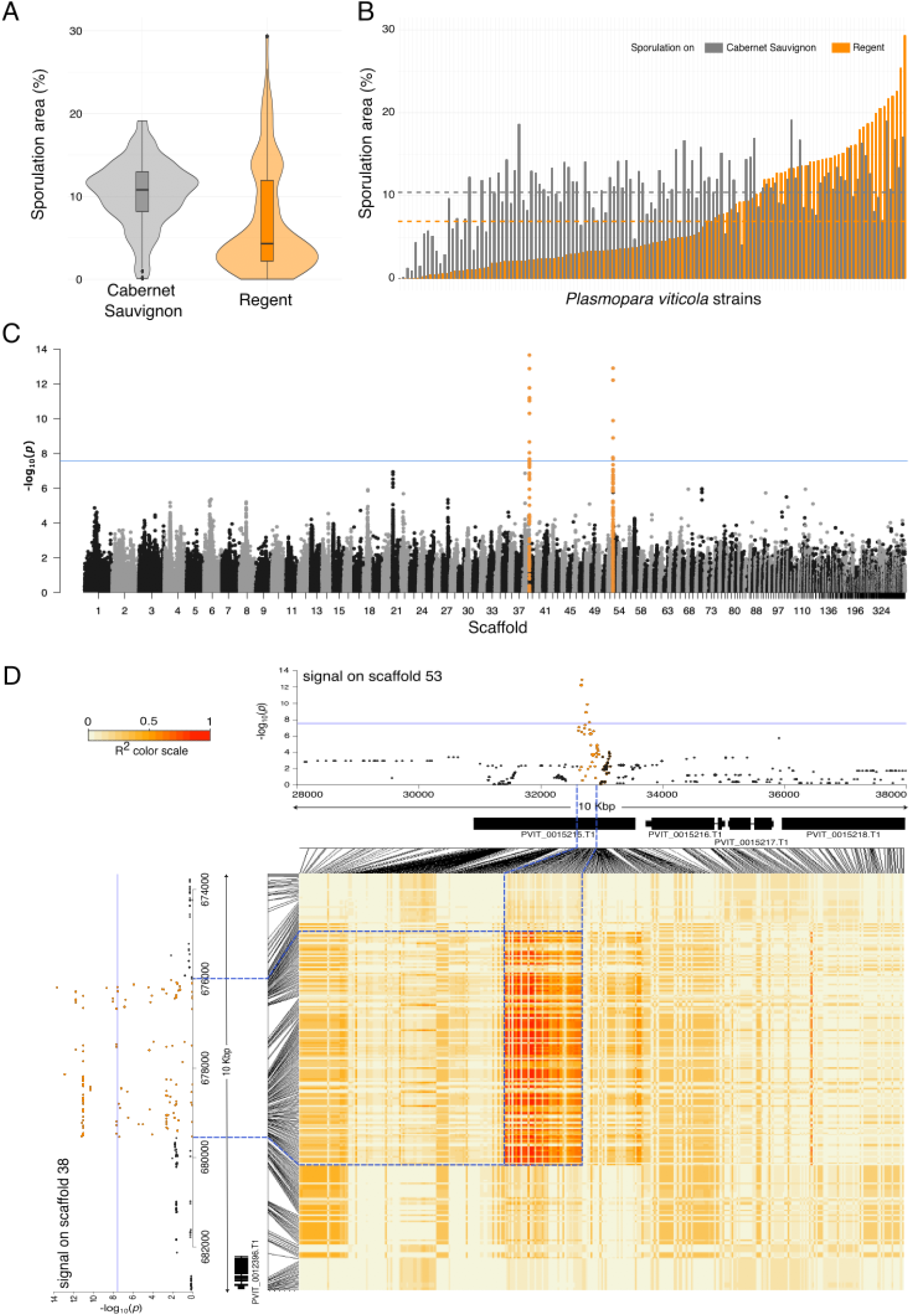
Result of association genomics analysis of *P. viticola* virulence on the resistant variety Regent carrying Rpv3.1. (A) Boxplots showing the distribution of the sporulation area of the population on Cabernet Sauvignon (Rpv3.1-, grey) and Regent (Rpv3.1+, orange). (B) Histogram of the sporulation area of each of the 123 strains on Cabernet Sauvignon (grey) and Regent (orange). Values correspond to the mean of sporulation area in percentage for six replicates. The horizontal dashed lines represent the average of the population on Cabernet Sauvignon (grey) and Regent (orange). The standard error and strain names are not represented for readability of the figure and are available in SI dataset S1. (C) Manhattan plot of the whole genome. Each point corresponds to a single SNP. The x-axis represents the physical location of each SNP on the genome (split by scaffold). The y-axis represents the significance of the association between a SNP and the studied phenotypic trait. The blue horizontal line indicates the significance threshold corrected by the Bonferonni criterion (alpha=0.5). The two loci significantly associated with virulence on Rpv3.1 are indicated in orange. (D) Linkage Disequilibrium (R^2^) between the two signals identified by GWAS. The heatmap illustrates the R^2^ values between each SNP in the signals from scaffold Plvit038 (y-axis) and SNPs in the signals from scaffold Plvit053 (x-axis). The two Manhattan plots provide a zoomed-in view of the two targeted loci located on scaffolds Plvit038 and Plvit053, as presented in panel C. The locations of genes within the locus are indicated below the Manhattan plots. Correspondence between SNPs in the Manhattan plots, their positions in genes, and their R^2^ values in the matrix is highlighted using blue dashed lines.

We used the sporulation area of the strains on Regent as a quantitative trait to perform a Genome-Wide Association Study (GWAS). We used a Genome wide Efficient Mixed Model Association (GEMMA) to correct for the population structure bias evidenced by the principal component analysis (Figure S2). The Quantile-Quantile (Q-Q) plot showed no evidence of inflation in test statistics for the sporulation on Regent (*λ*=1.04) (Figure S3). Finally, the proportion of phenotypic variation explained (PVE) estimated by GEMMA was 0.58, indicating that genetic variants account for 58% of the phenotypic variance in our population.

We identified a total of 74 markers strongly associated with this phenotypic trait (Figure 1C). On the scaffold Plvit038, the 66 markers are located in an interval of 3,340 bp (Figure 1D and S4 A and B) and exhibit strong linkage disequilibrium (Figure S4 D). No gene nor pseudogene was detected at this location on the genome assembly (Figure 1D and S4 C). In the locus on scaffold Plvit053, 8 markers, in an interval of 144 bp, were significantly associated with the sporulation on Regent (Figure 1D and S4 A and B) and are in strong linkage disequilibrium (Figure S4 D). The significant SNPs identified by GWAS are at the edges of Plvit038 and Plvit053 scaffolds (Figure S4 A) and show evidence of being tightly associated as observed by a strong linkage disequilibrium (Figure 1D). The high R^2^ values around 0.7 observed between the two scaffolds indicates a close physical proximity of the sequences, which appears to have been disrupted by the fragmentation of the genome assembly. Based on the gene annotation (19), one gene is located in this region: PVIT–0015215.T1 (Figure 1D).

### A structural rearrangement identified in the avrRpv3.1 locus

To discern any potential disparities between haplotypes in the haploid consensus reference genome and better understand the link between the two loci identified by GWAS, we performed a diploid-aware *de novo* assembly using FalconUnzip (53). This approach allowed us to reconstruct and annotate the gene content for both alternative haplotypes, as presented in Table S2. The primary assembly, which represents the most contiguous haploid representation of the genome, comprises 80.6 Mb divided into 252 primary contigs, containing 23,602 protein coding gene loci. The genome size of this new assembly (80.6 Mb) is comparable to previous assembly results but falls intermediate between a previous SMRT sequencing assembly (92.94 Mb, (19)) and an Illumina assembly (74.74 Mb, (54)). This is attributed to the combined effect of improved representation of repetitive content enabled by long reads and the utilization of a diploid-aware procedure capable of effectively segregating highly divergent alleles into haplotigs (i.e., phased alternative haplotypes). Moreover, with the longest sequence reaching 3.17 Mbp and an N50 of 825.8 kbp (N50 index 29), the new diploid assembly exhibits enhanced sequence contiguity compared to previous *P. viticola* strain INRA-Pv221 assemblies (19, 54). Falcon-Unzip reported a separate alternative haplotype representation for over 66.5% of the *P. viticola* genome (53.6 Mb in 745 Haplotigs, 15,642 protein coding gene loci), confirming the extensive structural variability present between haplotypes in the INRA-PV221 strain.

By aligning the scaffolds Plvit038 and Plvit058 with the sequences of both primary contigs and haplotigs of the diploid reference, we confirmed the contiguity of the two scaffolds within the same genomic region (Figure S5 A and B). This was ascertained by their juxtaposition on both a primary contig (Primary_000014F) and one of its associated haplotigs (Haplotig_000014F004), as illustrated in Figure S5 C and D. Additionally, we discovered the presence of structural variations between the two haplotypes within this locus, hereafter named avrRpv3.1. These structural variations were identified in the vicinity of the anticipated gene locus that encodes PVIT_0015215.T1. A major structural event involving a 30 kbp deletion was observed (Figure 2 and Figure S7) at this locus. Specifically, the genomic region spanning from 695 kbp to 725 kbp in Primary_000014F is absent from Haplotig_000014F004. This finding suggests that the consensus haploid assembly (19) exclusively represented the structure of the locus represented in the haplotig sequences. The deletion and the other structural variants found in the locus affects three coding gene loci: Primary_000014F.g163, Primary_000014F.g164 and Primary_000014F.g165 (hereafter called g163, g164 and g165). The gene g163 is translocated along with a transposable element (TE) found in this region (Figure 2). We discarded g163 from the following analyses as it is not deleted and is related to TEs. The genes g164 and g165, which were absent in the consensus assembly, are deleted in the haplotig and were not found to be duplicated in the genome. Gene PVIT_0015215.T1 of the consensus assembly corresponds to gene g166 from the diploid assembly (99.9% coverage and 98% identity), located right next to the deletion and present in both haplotypes (Figure 2). These genes are called hereafter avrRpv3.1 genes.

**Fig. 2.**
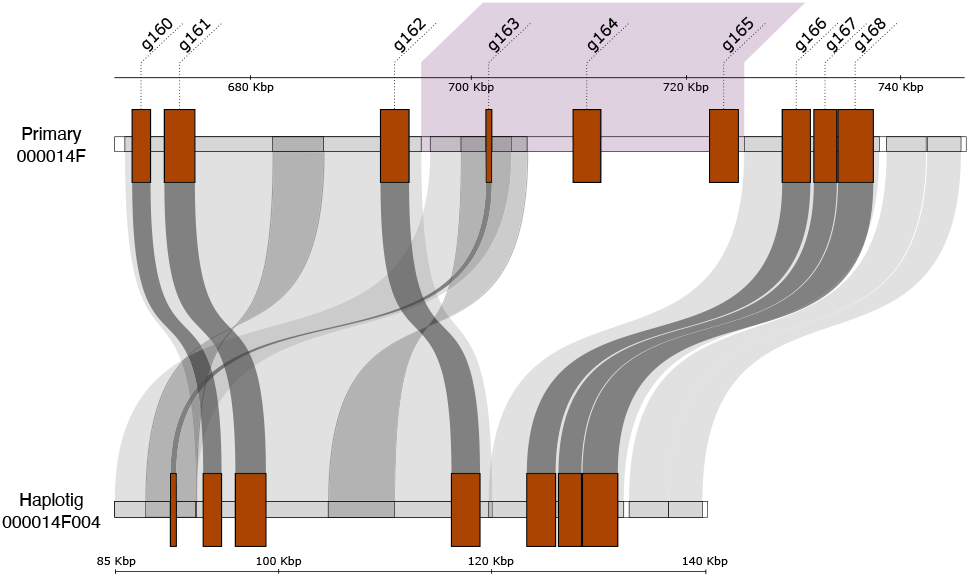
Structural representation of the locus associated with virulence against Rpv3.1 resistant grapevine, avrRpv3.1, on both haplotypes of the reference genome INRA-Pv221. The genes are shown as brown squares. The deletion associated with this locus, identified through GWAS, is highlighted in purple. The links between the Primary and Haplotig sequences (as well as between genes) connect regions showing significant homology, indicating that the sequences from the Primary are similar to those on the underlying Haplotig (Figure S6).

To gain a comprehensive understanding of the distribution of avrRpv3.1-like genes in *P. viticola* and other oomycete plant pathogens, we utilized the sequence of g164 as a query and conducted a search through oomycete genomes available in GenBank using BlastP. A total of 11 sequence matches were found in *P. viticola* genome and 14 sequences in the genome of *P. halstedii*, one of which being annotated as an RXLR-like protein. No sequence matches were observed outside the genus *Plasmopara*, including *Bremia, Phytophthora* and *Pythium* species. Interestingly, the 11 sequences of *P. viticola* were organized in a single cluster within the scaffold Primary_000014F (Figure S7). Within *P. viticola*, the gene size ranged from 2,535 to 2,688 bp with a mean size at 2,650 bp. The 11 *P. viticola* proteins presented a high degree of similarity estimated by pairwise comparison of sequences (average of 61% identity). The phylogenetic analyses (Figure S7) indicated that g162, g164, g165, g166 and g169 formed a well supported group among which g164 and g165, the two genes included in the deletion, where the closest relatives (69.76% of conserved amino acids). Altogether, the analyses of avrRpv3.1 candidate genes evidenced that a genus-specific gene family expansion has occurred through tandem duplication events in the *P. viticola* genome.

### Allelic diversity at the avrRpv3.1 locus

We explored the allelic variation at the avrRpv3.1 locus (interval between 640 and 740 kbp of Primary_000014F) in the 123 *P. viticola* strains. For each strain, we performed a diploid-aware copy number variation (CNV) analysis to identify regions that were significantly different in coverage in the locus from the rest of the contig (Primary_000014F) and detect the underlying structural variants (Figure 3A and Figure S8-S9). We observed that the deletion at the locus avrRpv3.1 is variable in size, evidencing the existence of multiple allelic forms. We identified the presence of one allele without any deletion (Avr) and six alleles (avr1 to avr6) presenting deletions ranging from 14 kbp to 96 kbp Figure S10). The high density of TEs in this region impact the accuracy of the deletion coordinates. The deletion impacted the presence of genes g164 and g165. The five avr alleles share a deletion of the gene g164, which is complete in all alleles, except for avr2 where g164 is only partially deleted. The gene g165 is present in avr3, partly deleted in avr2 and avr4 and fully deleted in avr1, avr5 and avr6 (Figure 3B and Figure S10).

**Fig. 3.**
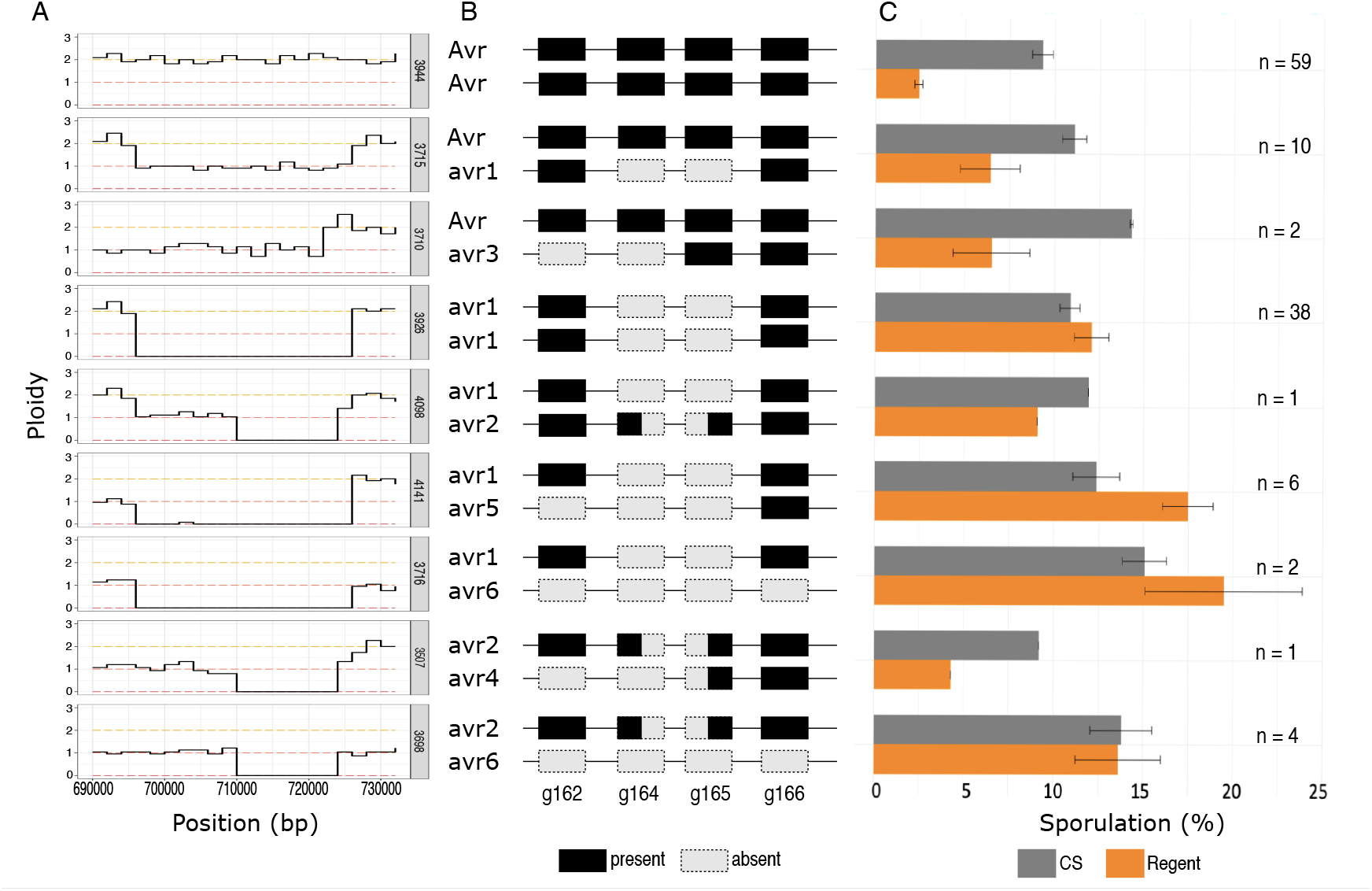
Allelic configuration analysis of the 123 *P. viticola* strains at the locus avrRpv3.1. (A) Patterns of coverage in the locus. A relative level of coverage of two indicates the presence of sequences both haplotypes, a level of one indicates that the sequence is present in only one of the two haplotypes, 0 copy means a complete deletion of the sequence in both haplotypes. Only one strain is represented per allelic configuration, details for all strains, from 640 to 740 Kpb, are available in Figures S8-S9. (B) Schematic representation of the locus in its diploid form with the genes (rectangles) involved. Black rectangles indicate the presence of the gene, grey rectangles indicate deletion of the gene, black and grey rectangles indicate a partial deletion of the gene. (C) Phenotype associated with each genotypes. Barplots indicate the sporulation area on Cabernet Sauvignon (CS) (grey) and on Regent (orange) observed for the isolates associated with to each of the allelic configurations. The number of strains of each category are indicated by *n*.

Through CNV analysis, nine distinct allelic configurations were identified among the 123 strains (Figure 3B): 59 were homozygous for the Avr allele, 12 were heterozygous (Avr/avr) and 52 display a deletion on both haplotypes (avr/avr). The Avr allele was thus the most frequent one (53%) followed by avr1 (39%). The frequency of the five other avr alleles represented 8% of the population and varied from 0,5% to 3%. The average sporulation area associated with each genotype on both Cabernet Sauvignon (CS) (Rpv3.1-) and Regent (Rpv3.1+) is depicted in Figure 3C. Strains with the Avr/Avr genotype, i.e., homozygous without any deletion, display four times more sporulation area on CS (mean = 9.31%) than on Regent (mean = 2.4%). The avr/avr genotypes have a similar or higher sporulation on Regent (mean = 12.91%) than on CS (mean = 11.42%). Finally, the Avr/avr genotypes display a sporulation two times more important on CS (mean = 11.62%) than on Regent (mean = 6.41%). It may be noted that strains Avr/avr have a sporulation level on Regent that is variable, ranging from 5% to 15% of the sporulation area (details for each strains displayed in Figure S12). Overall, we concluded that the breakdown of the Rpv3.1 resistance results from to the structural variations at the locus avrRpv3.1.

Additionally, we examined the avrRpv3.1 locus in another *P. viticola* strain collected from Switzerland that exhibited virulence on Rpv3.1-resistant varieties (strain avrRpv3-Rpv12-in (21)). Using the same pipeline analysis as for our *P. viticola* population, we observed a 30 kbp deletion in one haplotype and a 56 Kpb deletion in the other at this locus (Figure S12). This matched the avr1/avr5 genotypes of *P. viticola* identified in our population. Overall, the results obtained on *P. viticola* strains from a different geographical origin confirm that structural variations at the avrRpv3.1 locus are involved in the breakdown of Rpv3.1 resistance.

Based on the allelic configurations described above, the avrRpv3.1 locus revealed to be in strong Hardy Weinberg (HW) disequilibrium (p-value = 1.34e-21) with a negative D value (D = -24.65) suggesting a strong deficit of heterozygous genotypes. This result contrasts with results obtained along the contig Primary_000014F, where more than 95% of the SNPs were at HW equilibrium as expected for neutral markers in panmictic populations. We also observed a high genetic differentiation (F_ST_ = 0.58) between strains sampled on the susceptible and the resistant varieties at the locus while the average F_ST_ calculated across the contig was found to be 0.03 ± 0.05 with 99% of the F_ST_ values lower than the value calculated at the avrRpv3.1 locus (Figure S13). The heterozygote deficits and strong genetic differentiation between strains collected on susceptible and resistant grapevines may result from host selection occurring at the avrRpv3.1 locus.

### The candidate genes of the avrRpv3.1 locus encode putative effector proteins

The genes g164, g165 and g166 present in the region encoded proteins of 800-900 amino acids that were predicted to possess a signal peptide (SP) and that did not contain other motifs or domains based on InterPro searches. Alignment of the protein family revealed an EER motif, but absence of RXLR or RXLR-like motifs in their N-terminal sequence and the presence of repeated motifs (Figure S14). Structural similarity searches using HHPred and Phyre2 produced best hits with the structures of RXLRs effectors PSR2 from *Phytophthora sojae* and PcRXLR12 from *Phytophthora capsici*.

We next modeled the structure of g166 using AlphaFold2. The overall quality parameters of the model allowed to be confident in the backbone structure (Figure S15 A and B). The g166 predicted structure is horseshoe-shaped and similar to the Alphafold-predicted structure of a related protein from *P. halstedii* (Figure S15 C,D). Because the predicted structure appeared modular, we hypothesized that it was composed of repeats of the LWY-fold. We thus divided the g166 predicted structure into structural modules based on the PSR2 LWY domain structure, starting at the N-terminus with the alpha-helices corresponding to the helix bundle 1 (HB1) of the LWY domain. The g166 structural modules obtained, which we tentatively named LWY modules, produced a good overlap between them at the N-terminus but they were different at the C-terminus (Fig S16 A). Furthermore, superimposition of the PSR2 LWY motifs and the g166 LWY modules revealed an overlap at the level of the PSR2 LWY HB1 but a different structural organization for HB2 (Figure S16 B).

We divided the g166 predicted structure into modules that contained the HB1-like sequence at the C-terminal part, resulting in a protein consisting of 9 modular repeats, most containing 5 alpha-helices (Fig 4 A), which we tentatively named LW modules. Superimposition of g166 LW modules 2 to 9 revealed a good structural overlap (Fig 4 B), while LW module 1 appeared to be different. Superimposition of the HB1s from PSR2 and g166 LW modules showed a good overlap (Fig 4 C) and alignment of the primary sequences confirmed the conserved position of amino acids involved in maintaining the structure, with the exception of the PSR2 L2 position, which was replaced by a conserved Trp in g166 (Fig 4D). These observations suggest that the candidate genes found in the avrRpv3.1 locus encode putative effector proteins.

**Fig. 4.**
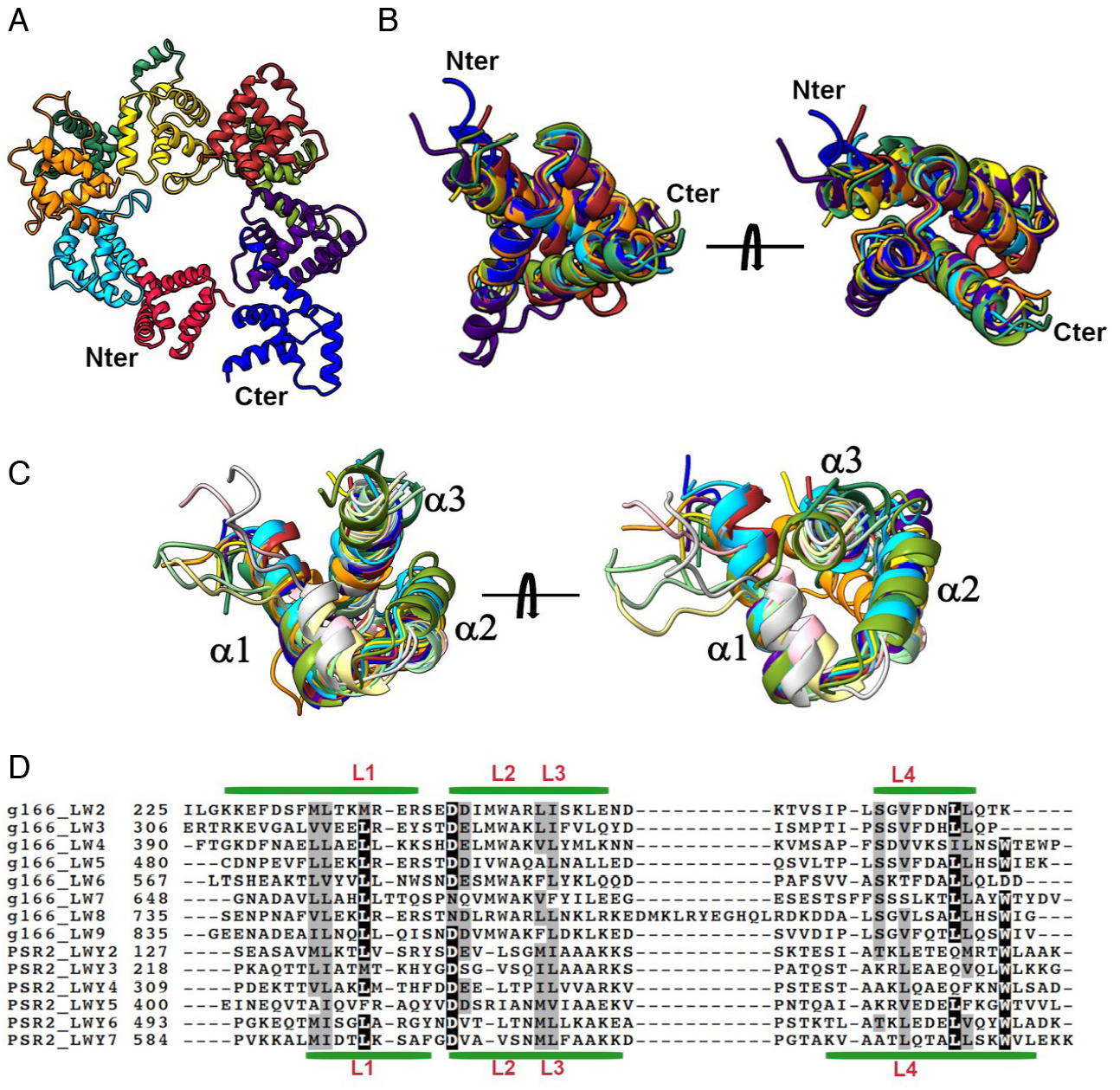
The predicted tertiary structure of g166 is composed by modules containing the HB1 fold from the LWY domain. (A) Predicted structure of g166. The first 108 amino acids have been removed for clarity. Modules 1 to 9 are shown with different coloring (1 to 9: red, light blue, orange, green, yellow, brown, olive, purple, blue). (B) Superimposition of LW modules 2 to 9 from g166, seen from two different angles. Coloring as in A. The N- and C-terminal parts of the modules have been trimmed for clarity. (C) Superimposition of the N-terminal helix-bundle 1 (HB1) from LWY-domains 2 to 7 from *P. sojae* PSR2 and the C-terminal HB1 from LW modules 2 to 9 of g166. Alpha-helices 1 to 3 are indicated. Coloring in g166 as in A. Coloring of PSR2 LWYs (2 to 7): light sky blue, khaki, sea green, silver, light pink, light green. (D) Alignment of the HB1 primary sequences from the g166 LW modules and the PSR2 LWYs. Green lines indicate alpha-helices 1 to 3 (from left to right) for g166_LW2 (top) and PSR2_LWY7 (bottom). Conserved Leu residues contribution to the HB1 fold are shown in red. Note the conservation of a Trp in the g166_LWs at the L2 position. Black background shows identity and grey background shows similarity (75 % cutoff).

### Candidate avrRpv3.1 proteins induce cell death in plants carrying Rpv3.1

Virulence towards Rpv3.1 was associated with the deletion of g164 and g165, making them the most promising candidates to be the cognate Rpv3.1 avirulence gene. We assessed the Rpv3.1-dependent cell death-inducing activity of avrRpv3.1 genes by transient expression of the proteins coded by g164, g165 and the closely related gene g166.

We confirmed the presence of the three genes in the avirulent strain Pv221 with a PCR on genomic DNA (Figure S17). RNA-seq data at 24-, 48- and 72-hours post-inoculation (hpi) revealed that g164, g165 and g166 showed to be modulated in a were expressed at very low level at 72 hpi (SI Dataset S6). The results were confirmed with a semi-quantitative RT-PCR revealing that all 3 genes are expressed in germinated spores and during infection (Figure S18). The expression pattern suggests weak constitutive expression.

We cloned the coding sequences for g164, g165, and g166, without the signal peptide, from the reference avirulent *P. viticola* strain INRA-Pv221. Two alleles were obtained for g166, which we named g166p and g166h according to the original haplotype. As shown in Figure 5, all genes induced cell death in Regent but not in Syrah. Induction of cell death by g166h was the stronger and more reproducible. Our results suggest that the candidate effector proteins from the avrRpv3.1 locus can induce Rpv3.1-dependent cell death.

**Fig. 5.**
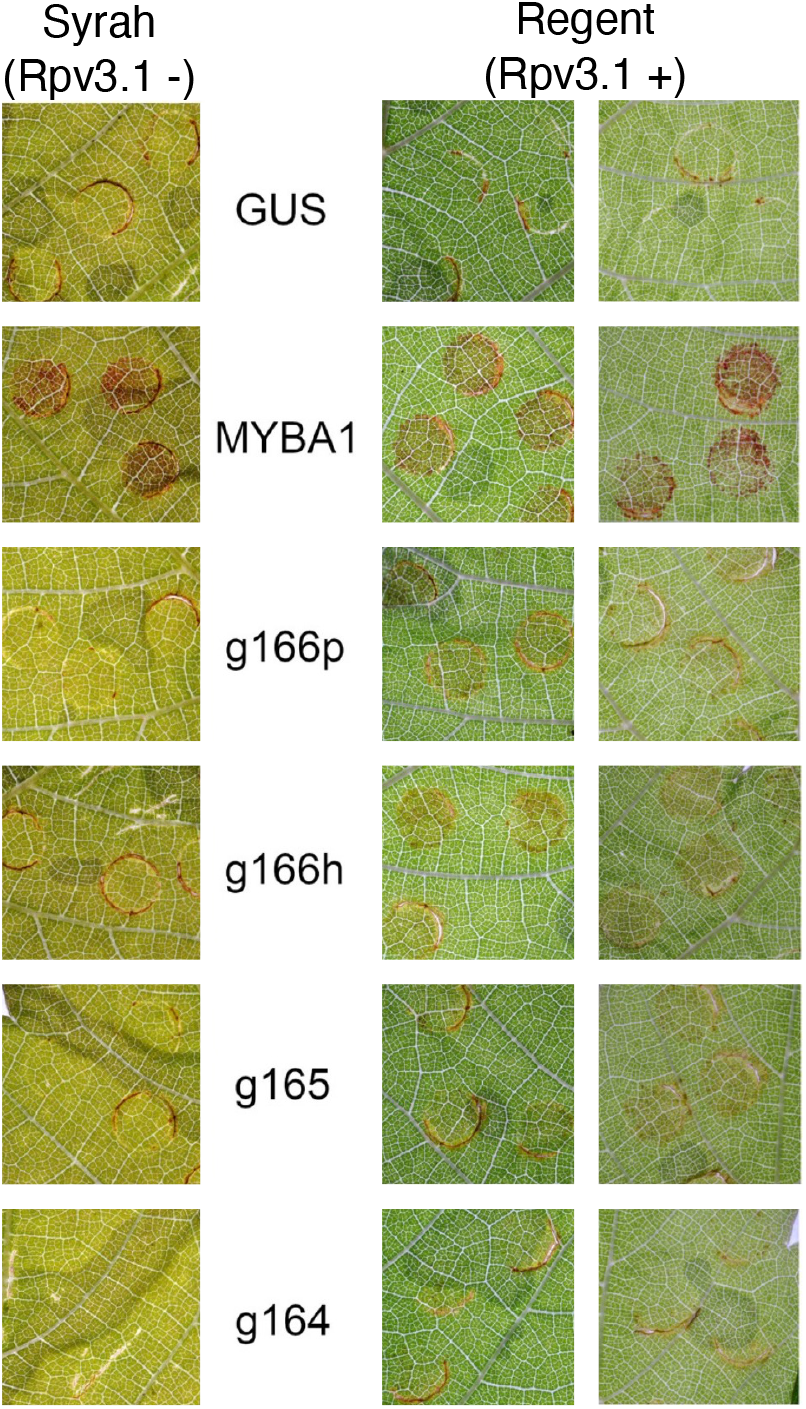
Candidate effector proteins from the avrRpv3 locus induce cell death in plants carrying Rpv3. Cell death induction in leaves from the grapevine varieties Regent (Rpv3.1+) and Syrah (Rpv3.1-) 6 days after Agrobacterium-mediated expression of effector candidate genes g166h, g166p, g165 and g164. Agrobacterium-mediated expression of *β*-glucuronidase (GUS) was used as negative control for unspecific induction of cell death following agroinfiltration. Agrobacterium-mediated expression of VvMYBA, leading to the production of anthocyanins, was used as positive control to assess the efficiency of transformation. For Regent, the result of two independent experiments out of the four conducted are reported.

## Discussion

By employing a GWAS approach, entailing the phenotyping of *P. viticola* strains and their whole-genome sequencing, we successfully identified the avrRpv3.1 locus involved in the interaction with the Rpv3.1-mediated resistance in grapevine. We presented compelling evidence, based on population genetics indices, that this locus displayed non-neutral characteristics and demonstrated signs of positive selection on resistant hosts. Within this locus, the deletion of two coding genes (g164, g165) is associated to the Rpv3.1 resistance break-down. These two genes are part of a cluster of 11 closely-related proteins, all exhibiting the characteristic traits of potential oomycete effectors, including the presence of a signal peptide and EER motif, as well as structural similarities to known RXLR effectors. Notably, these effector proteins possess an unusually large size for oomycetes, measuring approximately 880 amino acids. To put this into perspective, the largest known oomycete effectors to date are PsPSR2 (670 aa; (32)) and AVRcap1b (673 aa; (55)) and an analysis of the RXLR effector repertoire across seven oomycete species encompassing 2,126 proteins, revealed only eight proteins exceeding 800 aa in size (32). Through predicted structural modeling, we identified a modular arrangement within the candidate effector proteins, featuring repeats reminiscent of the LWY fold (32). Based on predicted structure, the identified modules overlapped with the LWY fold only at the level of the helix bundle 1 (HB1) suggesting a possible new fold in the effector repertoire of oomycetes. Prior to this research, no avirulence genes had been characterized in the oomycete pathogen *P. viticola*. This study therefore presents the first-ever documentation of avirulence genes responsible for breakdown of resistance in grapevine.

The experimental results showed that both candidate effectors (g164, g165) and the closely related gene (g166) induce cell death in presence of Rpv3.1, suggesting that the gene/s in the Rpv3.1 locus recognize all three effectors. The Rpv3.1 locus has been previously mapped to an interval containing two TIR-NB-LRR (TNL) genes, and it has been proposed that both genes are essential for conferring resistance (30). This raises the possibility that each TNL gene interacts with different effectors, potentially exhibiting varying degrees of recognition strength. Another aspect to consider is the introgression of Rpv3.1 in Regent, which involves a 15 Mb linkage drag encompassing a cluster of Nucleotide-binding and leucine-rich repeat (NLR) genes (56). It remains plausible that some of the candidate effector genes are recognized by other genes within this cluster. Overall, these findings suggest that Rpv3.1-mediated resistance may be the result of cumulative responses from the plant upon recognition of multiple effectors by one or more resistance genes. Consequently, the loss of some of these effectors by the pathogen could lead to the breakdown of resistance. We indeed observed that the deletion of genes g164 and g165 is strongly associated with a high sporulation area on Rpv3.1-resistant hosts by *P. viticola* strains. The deletion of avirulence genes serves as an effective evolutionary strategy employed by pathogens to evade recognition and overcome plant defenses. Similar mechanisms have been documented in various fungal plant pathogens, including *Leptosphaeria maculans* (57), *Zymoseptoria tritici* (46), and *Melampsora larici-populina* (58). In these cases, the loss of effector genes enhances the virulence of pathogens in Brassica crops, wheat, and poplar, respectively.

The presence of a multi-gene effector family accompanied by a high density of transposable elements (TEs) suggests genomic instability in the region. The rapid evolution of avirulence genes often occurs in genomic regions characterized by high TE content, punctual mutations, and structural variations (59, 60). Previous studies have shown that TE insertions near avirulence factors can have a significant impact on the virulence of fungal plant pathogens (49, 61, 62). Consistent with this notion, various structural variations such as deletions, inversions, and recombinations were detected at the avrRpv3.1 locus. Another aspect of the locus variability lies in the range of deletion sizes observed, ranging from 14 kbp to 96 kbp. TEs are likely to play a significant role in this genome evolution, as their mobility can expand the gene space through DNA duplication, interruption, or induction of effector gene deletion (60, 63, 64). These new insights may have important implications when choosing a strategy for deploying grapevine resistance genes. Theoretical studies highlighted how the evolutionary and epidemiological control provided by the main deployment strategies (mixture, mosaic, pyramiding), but also their relative ranking, crucially depends on a sound knowledge of the mutation rate leading to pathogen breakdown (see (65) for a review and (66) for an application to grapevine downy mildew). This study shows the diversity of the mutational events, such as the complete or partial gene deletion, leading to virulence. All these events contribute to greatly increase the overall mutation probability towards virulence (67). In this setting, the pyramiding strategy is not necessarily the most sustainable one for preserving the efficacy of grapevine resistance genes (66).

The genetic variability observed at the avrRpv3.1 locus in the *P. viticola* population provides interesting insights into the origin of avr alleles. Notably, we found that one avr allele, avr1, was highly predominant in virulent strains of the *P. viticola* population (80% of the population), including the vir3.1 strain collected in Switzerland (21). The presence of the avr1 allele in both Bordeaux (France) and Switzerland suggests the possibility of recurrent mutation at the avrRpv3.1 locus leading to the emergence of virulent strains. Alternatively, it is possible that the avr1 allele already preexisted at low frequencies in European populations of *P. viticola*, i.e. before the recent deployment of modern varieties carrying Rpv3.1. It should be noted that until the 1950s, French-American hybrids were extensively cultivated in Europe at large scale. It is therefore plausible that the avr1 allele, selected during that period, has persisted at low frequencies in *P. viticola* populations. Further investigations involving a more extensive survey across Europe would enhance our understanding of allele diversity at the avrRpv3.1 locus and the prevalence of these alleles in *P. viticola* populations.

In conclusion, our study confirms the effectiveness of genome-wide association studies (GWAS) in identifying genomic loci involved in both qualitative traits, as illustrated by the discovery of the mating type locus of grapevine downy mildew (20), and quantitative traits, as demonstrated by the description of the locus responsible for the breakdown of a partial resistance in this study. We have shown that the identification of effectors, typically found in rapidly evolving genomic regions, would not have been possible without the utilization of diploid-aware genome assemblies. Our study underscores the advantages of working with diploid-aware genome assemblies, particularly in highly heterozygous genomes like the one of *P. viticola* (19). Looking ahead, the availability of highly accurate long reads, such as HiFi reads (68, 69), will greatly enhance our ability to obtain diploid references at the chromosome level. This advancement holds significant promise for studying genomic evolution in diploid organisms like *P. viticola*. By leveraging such advanced techniques, we can unravel complex genomic traits and deepen our understanding of gene-for-gene interaction. While our current knowledge of the molecular interactions between downy mildew and its host is still in its early stages, our research contributes to expanding this knowledge and opens avenues for further exploration of gene-for-gene interaction in this pathosystem. The recent discoveries regarding the breakdown of the partial resistances Rpv10 and Rpv12 in grapevine (21, 22, 26) pave the way for future exploration of new effectors in *P. viticola* using similar approaches.

## Materials and Methods

### Strains and plant materials

We collected 136 *P. viticola* isolates in 2018 from two plots in Bordeaux vineyards located 5 km from one another (France). Further details about their origin are provided in SI Dataset The monosporangium isolation was carried out as described in (22). After monosporangium isolation, isolates are referred as “strains”. After two weeks of propagation (see supplementary materials), on the day of inoculation, the strains were suspended in sterile water and the density of the suspension was adjusted to 5 × 103 sporangia *ml*^*−*1^ in a volume of 120 ml. Two host plants were used for the phenotyping experiment. *Vitis vinifera* cv. Cabernet Sauvignon (Rpv3.1-) was chosen as a susceptible host to downy mildew and the variety ‘Regent’ as a partially resistant host carrying the resistance Rpv3.1 (Rpv3.1+). The plants, 75 of each variety, were grafted onto the SO4 rootstock and grown simultaneously in a glasshouse under natural photoperiod conditions, without chemical treatment. The cross-inoculation experiment was conducted on the fourth leaf below the apex (one leaf per plant) which was collected after six months of cultivation.

### Phenotyping experiment

We inoculated the 136 strains on the susceptible variety *V. vinifera* cv. Cabernet Sauvignon and the resistant variety ‘Regent’. Four ‘mock strains’, consisting of sterile water only, were used as negative controls. For each of the 280 interactions, we performed six replicates. We therefore generated 1680 Plant-Pathogen interactions in total (i.e. grapevine leaf disc inoculated with a *P. viticola* strain) as described in supplementary materials. The leaf-discs were placed in 36 square Petri dishes (23 × 23 cm), with each of the six replicates placed in a different Petri dish. Petri dishes were sealed with Parafilm to maintain relative humidity at 100%. They were then incubated in three growth chambers, paying attention to have the six replicates equally represented in the three different growth chamber, for six days at 18°C under a 12-hour light/12-hour dark photoperiod. Six days post inoculation, all leaf-discs were photographed and the sporulation area per disc determined by image analysis (code available at https://github.com/ManonPaineau/image_analysis_P.viticola) as described in (22).

### Genotyping and Genome-Wide Association Study

The DNA preparation, the sequencing, the variant calling, quality control of the SNP and the population genetic structure analysis were carried out as described in supplementary materials. The genome-wide association study was performed on the sporulation area quantitative trait using the exact Genome-wide Efficient Mixed Model Association (GEMMA) method (70). The association tests were realized with the relatedness matrix, estimated with the GEMMA software (http://www.xzlab.org/software.html), phenotype (average sporulation area value on Regent (Rpv3.1+) for the six replicates) and genotype (filtered SNPs). GEMMA also estimated the proportion of variance in the phenotype explained (PVE) by genotypes. We corrected the significance threshold by the Bonferonni criterion calculated as -log10(*α*/k)), where k is the number of statistical tests conducted and *α* = 0.05. We verified that the model fit our data and correctly accounted for population structure by checking the quantile-quantile plot and the degree of deviation of the genomic inflation factor lambda defined as the median of the resulting chi-squared test statistics divided by the expected median of the chi-squared distribution. The quantile–quantile plots and Manhattan plots were visualized with the R package qqman (71). We analyzed the linkage disequilibrium of the SNPs around the identified loci using the package LDheatmap implemented in R (72).

### Genome reassembly and avrRpv3.1 locus analysis

Whole genome assembly of Single Molecule Real-Time (SMRT) reads(19) was performed in a two step procedure using the customized FALCON-Unzip pipeline reported in (73) (https://github.com/andreaminio/FalconUnzip-DClab). Detail of the assembly procedure are described in supplementary materials. We compared the two INRA-Pv221 assemblies to identify, in the new diploid assembly, the location of the loci identified by GWAS in the consensus reference genome. We used NUCmer with the minimum cluster size c=65 and visualize the result with mummerplot from the MUMmer3 software (74). To evaluate the copy number variation in the locus, the short reads of each sample were aligned separately with bwa mem (version 0.7.17 (75)) on *P. viticola* strain INRA-PV221 FalconUnzip primary assembly and copy number were evaluated on the entire dataset with CNVkit (ver. CNVkit 0.9.9, (76)). To visualize the allelic count, per-base mapping coverage was calculated with samtools depth (version 1.10 (77)), mean coverage value was calculated on windows of 1 kbp in size and normalized on the sample diploid whole genome mean coverage.

### Gene expression analysis

We investigate the expression of the genes present in the locus AvrPv3.1 using the available RNA-seq data from (19). Briefly, the RNA were extracted from *Vitis vinifera* cv. Muscat Ottonel leave after 24, 48 and 72 hours post-inoculation (*HPI*). For each *HPI* conditions, three replicates were produced, for a total of nine RNA-seq samples (SRR accession list available in SI Dataset 5). We performed the trimming, the alignment and the quantification of the expression of the transcripts as described in supplementary materials. In order to differentiate reads from *P. viticola* and *V. vinifera*, the paired-end reads were aligned against both genomes of INRA-Pv221, newly assembled with Falcon Unzip in this study, and the first haplotype of *V. vinifera* cv. Cabernet Sauvignon genome v.1.1 (78). The number of reads before and after trimming, and mapping on both genomes are displayed in the SI Dataset 5.

### Protein functional analysis

BlastP searches were performed at NCBI against the nr protein database. Signal peptides were predicted with SignalPv6.0 (79). Structural similarity searches were performed on Phyre2 (80) and HHPred (81), and displayed with Boxshade. Structural prediction of g166 structure was performed using Alphafold2 (82) implemented at ColabFold (83) using default settings. Visualization and superimposition of predicted structures was performed on UCSF Chimera X 1.5 (84). The predicted structure of the *P. halstedii* protein was retrieved from the EBI Al-phafold2 database (https://alphafold.ebi.ac.uk/entry/A0A0P1AQR5). RNA extraction, cDNA synthesis and PCR were done as in (85). Each sample from infected tissues consisted of 4 leaf discs. Briefly, after RNA extraction, DNAse treatment was done using the Invitrogen-Turbo DNA free kit, and first strand cDNA was synthetized using the RevertAid First Strand cDNA synthesis kit (Thermo Scientific). PCR amplifications consisted of 30 cycles of 20 s at 94°C, 20 s at 58°C and 60 s at 72°C, followed by a final extension step of 10 min at 72°C. Primers are listed in SI Dataset 4.

### Transient assay

The coding sequences of g164, g165 and g166 without the predicted signal peptides were PCR-amplified with Phusion polymerase (NEB) from genomic DNA of *P. viticola* strain Pv221-INRA. Following amplification with primers containing restriction sites, the PCR products were digested (NEB restriction enzymes) and cloned directionally into plasmid pBIN61. Plasmids were transformed into *E. coli* strain DH10B by electroporation. Qiagen Plasmid Mini Kit was used to extract the plasmids from the *E. coli* cultures, Agrobacterium strain C58C1 carrying the pCH32 plasmid was then transformed by electroporation. Identity of the clones was confirmed by sequencing. Primers used for cloning are listed in SI Dataset 4. Agrobacterium cultures were grown at 28°C in 5 mL of L medium containing kanamycin (50 *μ*g/mL) and tetracycline (2.5 *μ*g/mL). After 2 days, 1 mL of the bacterial suspension was used to inoculate 5 mL of L medium containing kanamycin (50 *μ*g/mL), tetracycline (2.5 *μ*g/mL), 10 mM MES, 150 *μ*M acetosyringone, and grown in the same conditions for one day. Bacterial suspensions were centrifuged and the pellets were resuspended in a solution containing 10 mM MES, 10 mM MgCl2 and 150 *μ*M acetosyringone. After 2–3 hours of incubation at room temperature, bacterial suspensions were infiltrated at an optical density at 600 nm (OD600) of 0.4 using a needleless syringe.

### Data availability

The sample sequences of *P. viticola* strains were submitted to NCBI SRA with project numbers PRJNA875296. PacBio SMRT reads produced for INRA-PV221 genome assembly were released under the project number PRJNA329579. RNAseq data were released under the project number PRJNA329579.

## ACKNOWLEDGEMENTS

The authors acknowledge the financial support of the French National Research Agency (ANR) under the grant 20-PCPA-0010 (PPR VITAE), the Comité Interprofessionel des Vins de Bordeaux, and the French government in the framework of the IdEX Bordeaux University “Investments for the Future” program GPR Bordeaux Plant Sciences. We also would like to thank Pr. Jochen Bogs, Dr Chantal Wingerter and Dr Birgit Eisenmann (Winecampus Neustadt, Germany) for kindly sharing with us the *P. viticola* strains characterized in Wingerter et al. 2021.

