## Supplementary materials for "An effector deletion leads to the breakdown of partial grapevine resistance to downy mildew"

### Supplementary Note 1: An effector deletion leads to the breakdown of partial grapevine resistance to downy mildew

#### Extended materials and methods

***Plasmopara viticola* propagation..** Each isolate consisted of a single sporulating lesion collected from a single infected grape leaf. The leaf fragments were rinsed with sterile water and left overnight in the dark to allow sporulation to occur. Fresh sporangia were collected and stored in liquid nitrogen for monosporangium isolation. The monosporangium isolation was carried out as described in (22). After monosporangium isolation, isolates are referred hereafter as "strains". Two weeks before the phenotyping experiment, the strains were propagated on five Cabernet Sauvignon detached leaf. After one week of incubation, they were propagated on four detached leaves for the phenotyping experiment. One day before the experiment, the sporulating leaves were gently rinsed with distilled water to remove the sporangia already present, to ensure the collection of fresh sporangia of the same age on the following day. On the day of inoculation, the strains were suspended in sterile water and the density of the suspension was adjusted to 5 x $10^3$ sporangia $ml^{-1}$ in a volume of 120 ml.

***Plasmopara viticola* inoculation..** We performed the inoculation with a flotation method. For each strain, the 120 ml sporangia solution was poured into a sterile container and the 12 leaf discs (six replicates, two hosts) were placed on the surface of the sporangia solution, abaxial side facing the water. We used compartmentalized containers to avoid mixing the leaf discs while leaving the zoospores free to move throughout all compartments. When placing the discs on the solution, we ensured that the whole disc was in contact with the water thus avoiding air bubbles between the leaf surface and the water surface. The discs were left to float for three hours to allow the zoospores released into the solution from the sporangia to enter the leaf through stomata. After three hours of flotation, the discs were removed from the sporangia solution. After been carefully dried with sterile absorbant paper, discs were placed in 36 square Petri dishes for incubation.

**DNA extraction..** The DNA from the 136 strains was extracted directly from the mycelium of *P. viticola* with a DNA extraction protocol adapted from Möller et al. (1992), to which we added an RNAse A step. Briefly, the strains were firstly propagated on 4 Cabernet Sauvignon leaves. The mycelium was gently collected with a brush and then placed in a 2 ml tube containing TE buffer (10 mM Tris–HCl,1 mM EDTA). The suspension was centrifuged (20,000g,4°C), the supernatant was removed and the pellet was stored at -80°C. The samples were then lyophilized and grinded with 2 glass beads in a tissu lyser II Retsh grinder at 30s/sec. for 1 min. A volume of 500µl of TES (100 mM Tris,pH8.0; 10 mM EDTA; 2% SDS) lysis buffer and 10 µl Proteinase K (20mg/ml) was added for each sample and left to incubate for 30 minutes to 1h at 65°C in a water bath with an occasional gentle mixing. We then added 140µl of NaCl (5M) and 65µl of CTAB (10%) before another 10 minute incubation in a water bath at 65°C. The samples were gently vortexed between each step. We added 70µl of RNase A and then gently homogenized the tubes by inversion and let them rest for 30 minutes at 37°C. A volume (about 785µl) of chloroform/isoamylalcool (24:1) was added and gently agitated by inversion. After 30 min at 0°C, the samples were centrifuged for 10 min at 14000 rpm at 4°C. The supernatant was collected in a new tube. After gently stirring, we added 1/10 volume of NH4Ac (7.5M), and 1 volume of glacial isopropanol for an overnight incubation at -20°C. The next day, after 30 min of centrifugation at 14000 rpm at 4°C, the supernatant was removed to keep only the DNA pellet. A volume of 500µl of glacial ethanol (70%) was added for the washing step and the tubes were centrifuged again 10 min at 8000 rmp at 4°C. Finally, the tubes were emptied to leave only the DNA at the bottom of the tube. After drying, the DNA was suspended in 50µl of TE, left for 5 min at 37°C and stored at -20°C until sequencing.

**Sequencing..** The DNA quantity and purity were determined using a a Qubit fluorometer (Invitrogen, Carlsbad, CA,USA) and a Denovix (DS-11, DeNovix Inc., USA). DNA-seq libraries were constructed with the Illumina TruSeq nano kit. Sequencing was performed at the GeT-PlaGe facility (Toulouse, France) with a NovaSeq6000 and a Hiseq4000 (2x150 bp paired-end reads), except for the Pv221 individual, for which sequencing reads were already available (SRA accession numbers: SRX1970160, SRX1970161, SRX1970162) (19, 54).

**Variant Calling..** We first practiced a quality-check with FastQC analysis on each fastq paired read for quality control. Then a nextflow pipeline (https://forgemia.inra.fr/fabrice.legeai/pipelines/-/tree/master/Nextflow/SNPCalling) GATK_haplotypeCaller.nf was used to parallelize the GATK tools (version 4.1.4.1 (87)). After a fastp step for cleaning, mapping against INRA-Pv221 reference genome was done with bwa mem (version 0.7.17 (75)) and merging of differents bam files per library was done with samtools merge (version 1.10 (77)). Then, a GATK markduplicates step for the PCR duplication removal followed. Genotyping was then performed with GATK GenotypeGVCFs (Genomic Variant Call Format) with parameters output_mode EMIT_ALL_SITES and ERC GVCF. The average heterozygoty used by haplotypeCaller was adjust to het_mean = 0.001 and the standard deviation of heterozygoty to het_sd = 0.01. Join of gvcf files was made by another GATK_jointGenotyping_paral.nf pipeline which uses picard tools (http://broadinstitute.github.io/picard/), to CreateSequenceDictionary and gatk GenomicsDBImport for each intervals (equivalent to scaffold) before joining each gvcf with gatk GatherVcfs to obtain one cohort.vcf.

**Quality control..** We filtered out the variant sites present in repetitive regions with bedtools intersect (version 2.29.0 (88)) and then hard-filtered them using GATK recommendation thresholds (version 4.2.0.0 (87)). Variant sites with a minimum allele frequency (MAF) below 0.05 and with more than 10% of missing data in the population were excluded with vcftools (version 0.1.15 (? )). Sites with a minimum depth of 5 and a maximum of 3 times the individual mean coverage were considered valid calls using a homemade Python script. Finally, surviving sites with alternative calls not supported by at least five samples were removed. After filtering, the final dataset consisted of 123 strains for the GWAS panel encompassing a total of 1,854,765 polymorphic nucleotides.

**Population structure..** Population genetic structure was investigated by principal component analysis (PCA) with TASSEL (version 5 (89)). We subsetted the SNP dataset by selecting only the variants without missing data (with vcftools) and in low linkage disequilibrium by prunning them on a sliding window of 500 pb and a $R^2$ fixed at 0.2 with bcftools +prune (version 1.9 (90)). Population genetic structure was then assessed on a subset of 18,069 SNPs. The graphical representation of the PCA was carried out with ggplot2 (? ) implemented in R.

**Genome reassembly procedure and annotation.** Whole genome assembly of Single Molecule Real-Time (SMRT) reads(19) was performed in a two step procedure using the customized FALCON-Unzip pipeline reported in (? ) (https://github.com/andreaminio/FalconUnzip-DClab). Briefly, DAmasker (91) is applied to both raw and error corrected reads during FALCON-Unzip procedures (v.2017.06.28-18.01 (53)) to reduce the negative impact of repetitive content on the assembly performance in terms of assembly time and sequence contiguity. The first step was performed to optimize parameters for least fragmentation. Best assembly was obtained by using the following parameters: length_cutoff = 3000; falcon_sense_skip_contained = TRUE; falcon_sense_option = –output_multi –min_idt 0.70 –min_cov 4 –max_n_read 400; length_cutoff_pr = 15000; pa_DBsplit_option = -x500; pa_HPCdaligner_option = -v -dal128 -t30 -e0.7 -M40 -l1000 -k16 -h64 -w7 -s1000 -mtan -mrep2 -T4; ovlp_DBsplit_option = -x500; ovlp_HPCdaligner_option = -v -B128 -M40 -t60 -k20 -h256 -e.96 -l2000 -s100 -mtan -mrep2 -T8 ; overlap_filtering_setting = –max_diff 100 –max_cov 400 –min_cov 3.
After the Falcon procedure, assembled sequences were then compared with the RefSeq genomes database (retrieval date) using BLAST (ver. 2.2.28+, (92)) in order to identify potential contaminants. All sequences that best matched with non fungal genomes for over 50% of the length were considered as putative contaminants. Reads related to the putative contaminant contigs were removed from the successive assembly procedures.
Falcon assembly was then performed again using only the filtered SMRT reads dataset, followed by Unzip with default parameters and polishing with Arrow to remove residual sequence errors. The resulting draft assembly consisted of 253 Primary contigs delivering a contiguous haploid representation of *P. viticola* isolate INRA-Pv221 covering 80.4Mb. Over 66.5% of the genome was phased in a diploid representation with the alternative alleles assembled in 745 Haplotigs for a total of 53.5Mb. Draft sequences were searched for repeats using RepeatMasker (ver. 4.0.6, peronosporales database (93)). Gene prediction was obtained using Braker (ver. 1.9, (94)) trained on known *P. viticola* genes. CDS sequences were extracted for genes annotated in *P. viticola* in (54) and aligned on the diploid draft assembly using GMAP (ver. 2019-09-12 (95), models detected from alignments with identity and coverage greater than 95% and defining a complete ORF were used as hints in Braker for training. Prediction was further refined by realigning on the genome all the predicted gene models using GMAP (ver. 2019-09-12, (95)) and BLAT (ver 36x2, (96)). Matches with identity and coverage greater that 70% were used to identify previously unannotated gene coding loci and putative pseudogenes. The procedure allowed to identify 23,602 protein coding genes loci on the primary sequences and 15,642 protein coding genes loci in the alternative haplotypes.

**Phylogenetic analyses of the avrRpv3.1-like gene sequences.** To identify the avrRpv3.1-like gene sequences, the candidate effector g164 was used as query to search the *P. viticola* phased genome and all oomycetes genomes available via tBlastn with an E-value cut-off of e-10. The nucleotide sequences identified were aligned with Muscle (97), and the phylogenetic relationships were inferred by PhyML using the GTR model of nucleotides substitution (98). The trees were visualized with FigTree (99).

**Population genetic analysis.** In order to evaluate the influence of evolutionary forces on the locus avrRpv3.1, $F_{ST}$ (100) was measured between strains sampled on the susceptible variety and strains sampled on the resistant varieties at this locus and compared to the scaffold-wide distribution. The short reads of each *P. viticola* strains were mapped against the contig Primary_000014F (i.e. the contig containing the locus avrRpv3.1) of the newly assembled falconUnzip genome of INRA-Pv221 following the mapping, calling and filtration procedure described above. After filtration, a total of 29,443 SNPs were obtained and used to compute the $F_{ST}$ for each SNP distributed on the contig Primary_000014F with vcftools v0.1.16 (101). $F_{ST}$ was also computed specifically for the locus avrRpv3.1 using the "hierfstat" R-package (102). Additionally, Hardy-Weinberg Equilibrium (HWE) was tested for each SNP distributed on the scaffold with vcftools v0.1.16 (101) and specifically for the avrRpv3.1 locus using the "HardyWeinberg" R-package (103). This analysis aimed to identify any potential deviations from HWE, which could suggest a biased distribution of alleles at this locus resulting from selection pressures.

**avrRpv3.1 locus analysis of the virulent strain avrRpv12-/3-..** We obtained the *P. viticola* strain named "avrRpv12-/3-" sampled in Switzerland and demonstrated to be virulent on four resistant varieties carrying the Rpv3 gene (21). In the present study, we sequenced the DNA and described the locus avrRpv3.1 in this new strain, in order to investigate whether this locus is involved in bypassing Rpv3 resistance in a different *P. viticola* population. DNA extraction, sequencing, variant calling, quality controls and copy number variation analysis were carried out in the same way as for the other strains, following the protocol described above.

**Gene expression analysis..** We performed the trimming, the alignement and the quantification of the expression of the transcripts using, respectively, Trimmomatic v.0.36 (104) (settings: LEADING:7 TRAILING:7 SLIDINGWINDOW:10:20 MINLEN:36), the splice aware aligner HiSat2 v.2-2.1.0 (105) (settings: –non-deterministic –very-sensitive -t) and Salmon v.1.5.1 (106) (settings: -l U –numBootstraps 100 –seqBias –posBias ).

#### supplementary figures and tables

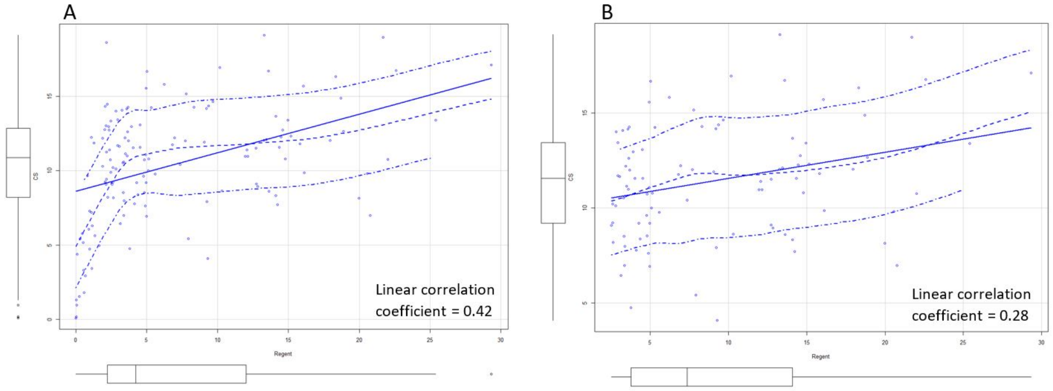

**Fig. S1.** Linear correlation between the sporulation area on Cabernet Sauvignon (CS) in y-axis and the sporulation area on Regent on the x-axis indicating that a strain characterized by a high sporulation on Regent is not characterized by a high sporulation on Cabernet Sauvignon. (A) A weak sporulation on Regent (less than 2.5%) is associated with low sporulation on Cabernet Sauvignon. For a higher sporulation value (greater than 5%), high sporulation on Regent appears to be indistinctly associated with both high and low sporulation values on Cabernet Sauvignon. The Pearson linear correlation coefficient is 0.42. (B) Without taking into account the individuals with a weak level of sporulation (< 2.5%) on Regent, we obtain a low Pearson linear correlation coefficient of 0.28. The analysis has been conducted on 1620 samples (mock inoculation removed from the analysis as well as two damaged discs). We used the Pearson correlation coefficient to calculate the correlation between the sporulation measured on Regent and CS.

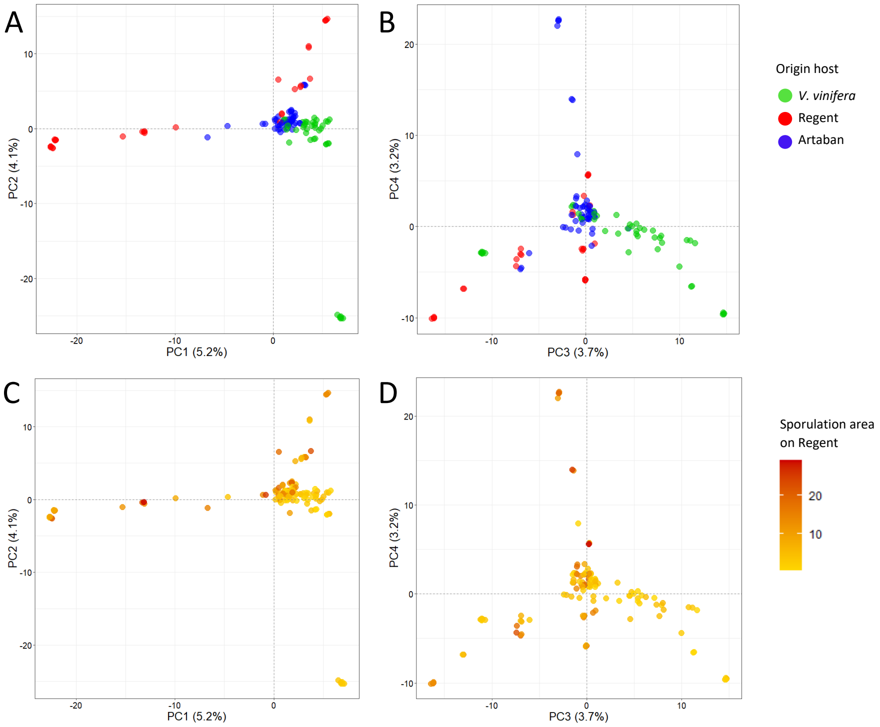

**Fig. S2.** Principal component analysis realized with a subset of 18,069 SNPs. A) and C) represent PC1 against PC2. B) and C) represents PC3 against PC4. A) and B) Strains are color coded according to their origin host. C) and D) Strains are color coded according to their sporulation area on Regent.

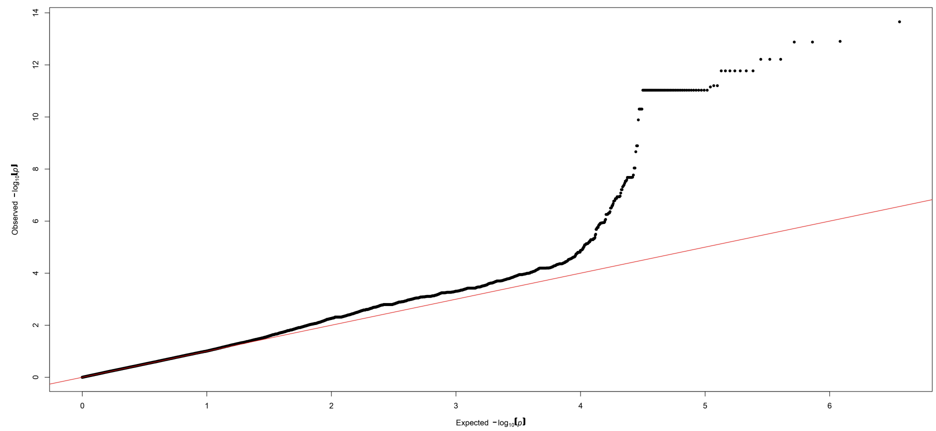

**Fig. S3.** Quantile-quantile plot associated with **Figure 1**. The distribution of markers (black dots) is close to the y = x line (red) as the observed -log10(P) are consistent with the expected -log10(P) for most of the markers. The deviation in the upper right tail of the y = x line suggests that an association is present in the data.

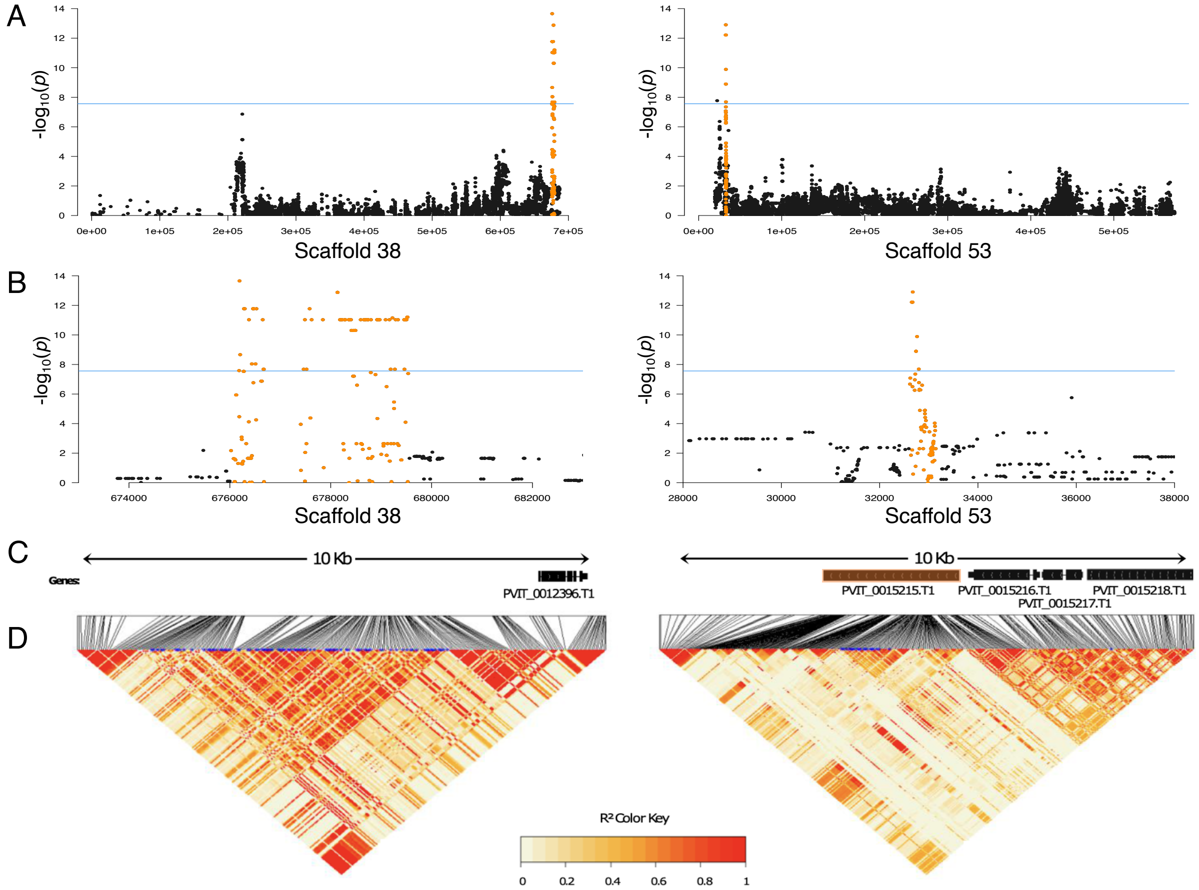

**Fig. S4.** (A) Manhattan plot of scaffolds Plvit038 and Plvit053 where the significant loci are located. The whole scaffolds are represented. (B) Zoom in for the two targeted loci located on scaffolds Plvit038 and Plvit053. A window of 10 kbp is represented. (C) The location of the genes on the loci. The gene of interest is highlighted in orange. (D) Heatmap of the Linkage Desiquilibrium ($R^2$) of the SNPs surrounding the two target loci shown in B). SNPs highlighted in blue correspond to the orange SNP from A and B.

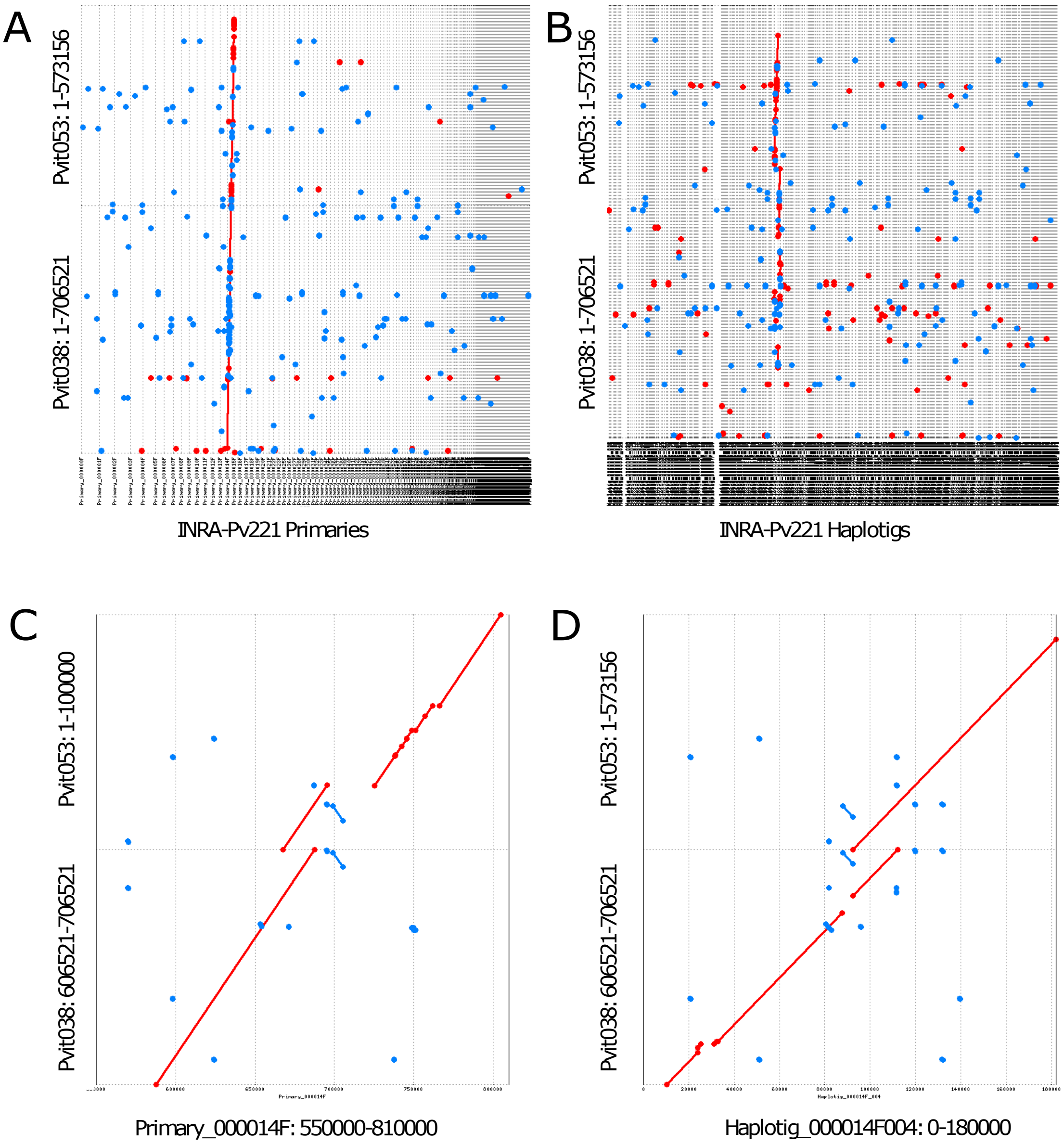

**Fig. S5.** Alignment of the sequences of Plvit038 and Plvit053 against INRA-Pv221 reference genome assembled with Falcon Unzip. Blue dots represent local forward alignments, red dots represent local reverse alignments. (A) Both Plvit038 and Plvit053 scaffolds are aligned on the contig Primary_000014F. All Primary contigs of INRA-Pv221 are used as reference. (B) Both Plvit038 and Plvit053 scaffolds are aligned on the Haplotig_000014F sequences. All Haplotig sequences of INRA-Pv221 are used as reference. (C) Alignment of the edges of the two scaffolds Plvit038 (the 10,000 last base pairs) and Plvit053 (the 10,000 first base pairs) on the Primary_000014F used as reference and (D) on the Haplotig_000014F_004.

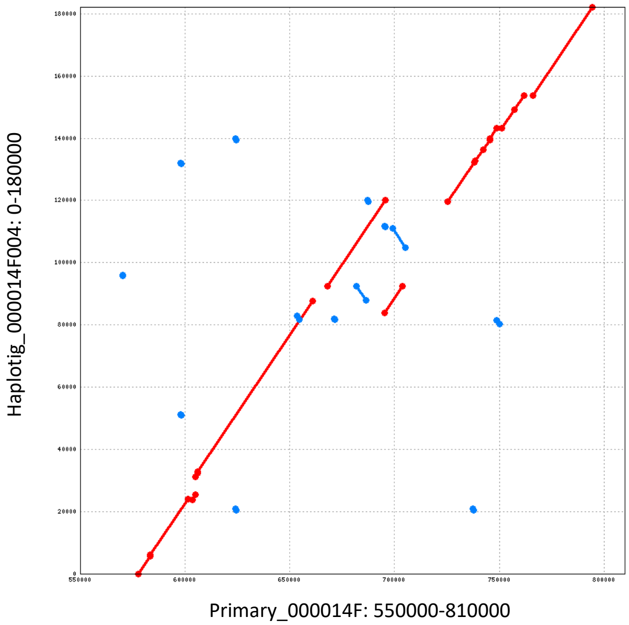

**Fig. S6.** Alignment of the sequences of the Haplotig_000014F_004 (the 180,000 first base pairs) against the Primary_000014F focused on the region of interest (between 550,000 and 810,000 bp). Blue dots represent local forward alignments, red dots represent local reverse alignments. The alignment coordinates serve as the basis for visualizing the locus in Figure 2.

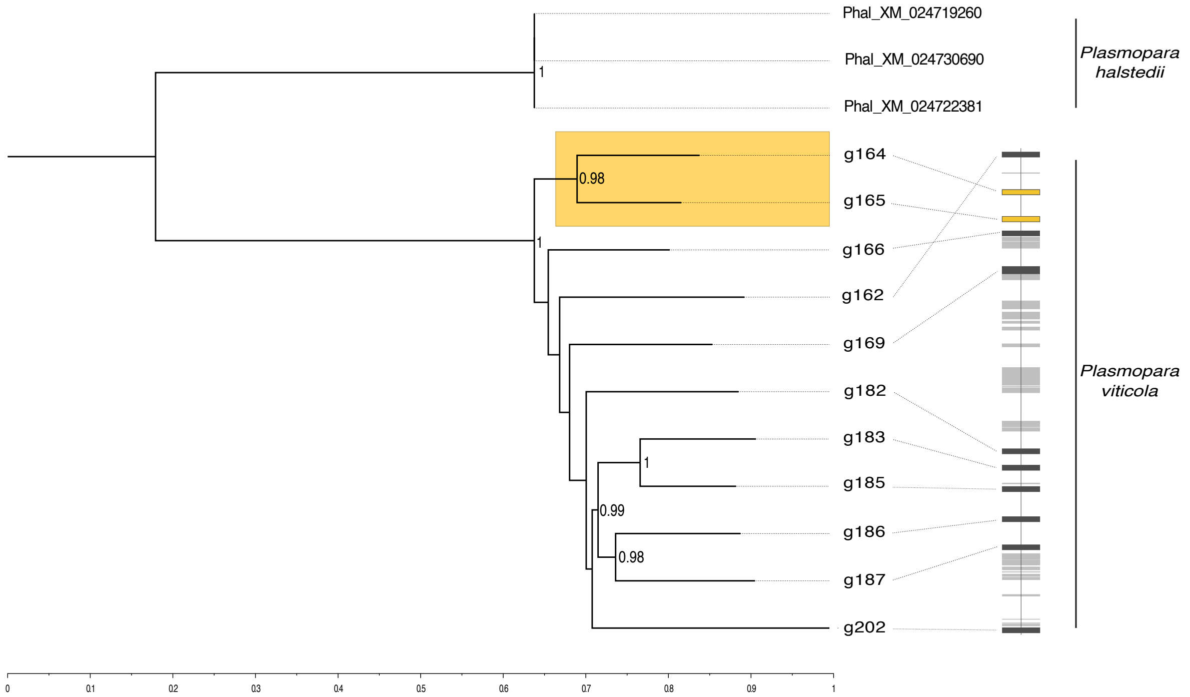

**Fig. S7.** Maximum likelihood phylogenetic tree of the 11 protein sequences of *Plasmopara viticola* identified as avrRpv3.1-like genes and three similar proteins found in the closely-related species *Plasmopara hasltedii*. Values of aLRT statistics > 50 are reported on nodes of the tree. Highlighted in yellow are candidate avirulence genes g164 and g165 that are deleted at the avrRpv3.1 locus in virulent *P. viticola* strains. The scaled gene track on the right showcases the physical position of the genes in the contig Primary_000014F of *P. viticola* genome. Dark grey denote avrRpv3.1-like genes (all other genes of the contig are in light grey).

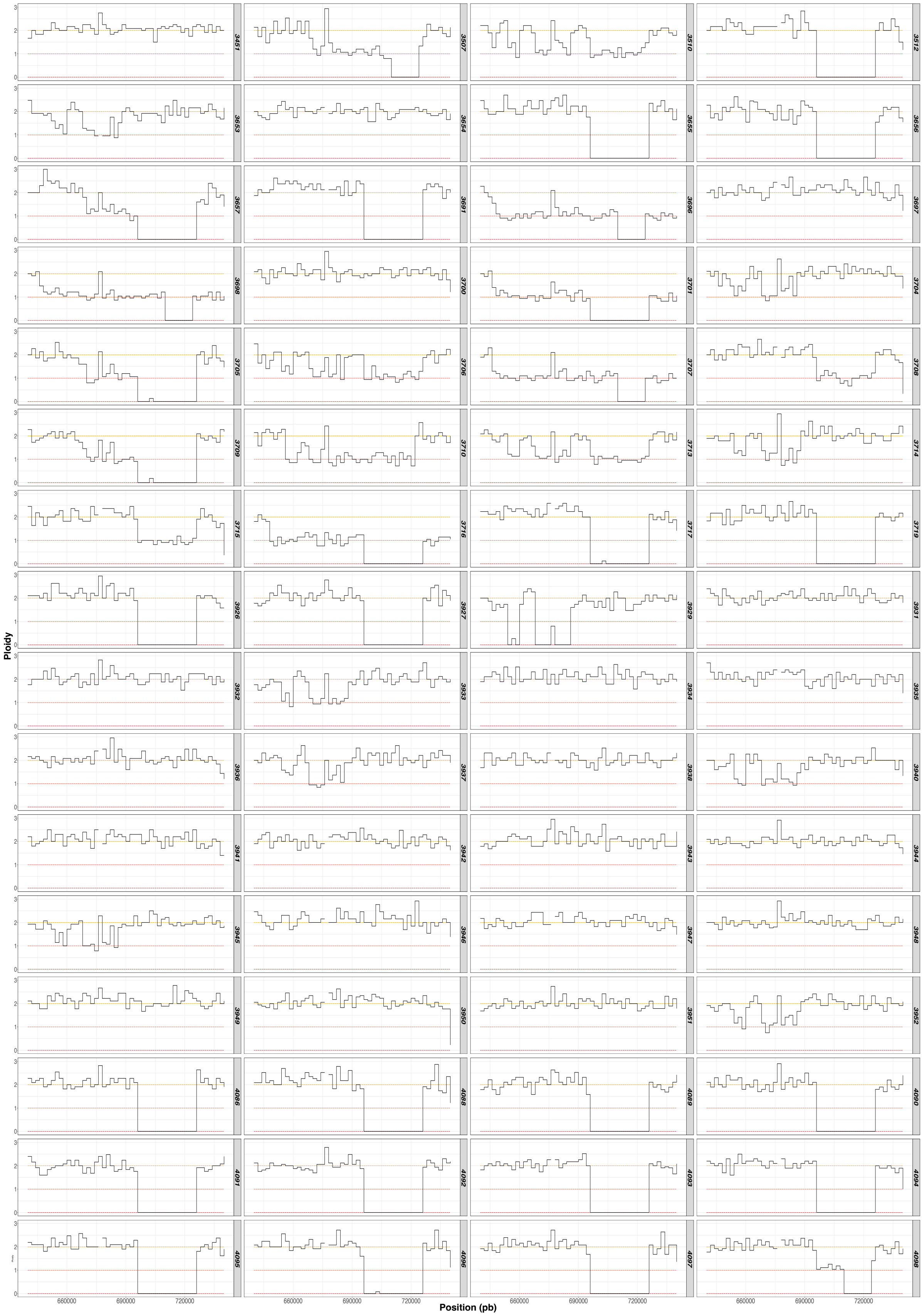

**Fig. S8.** Patterns of coverage in the locus (1/2). The analysis of the locus sequencing coverage between 640 Kpb and 740 Kpb unveils the patterns of deletions impacting the genes g164 and g165. A relative level of coverage of two indicates the presence of sequences both haplotypes, a level of one indicates that the sequence is present in only one of the two haplotypes, 0 copy means a complete deletion of the sequence in both haplotypes.

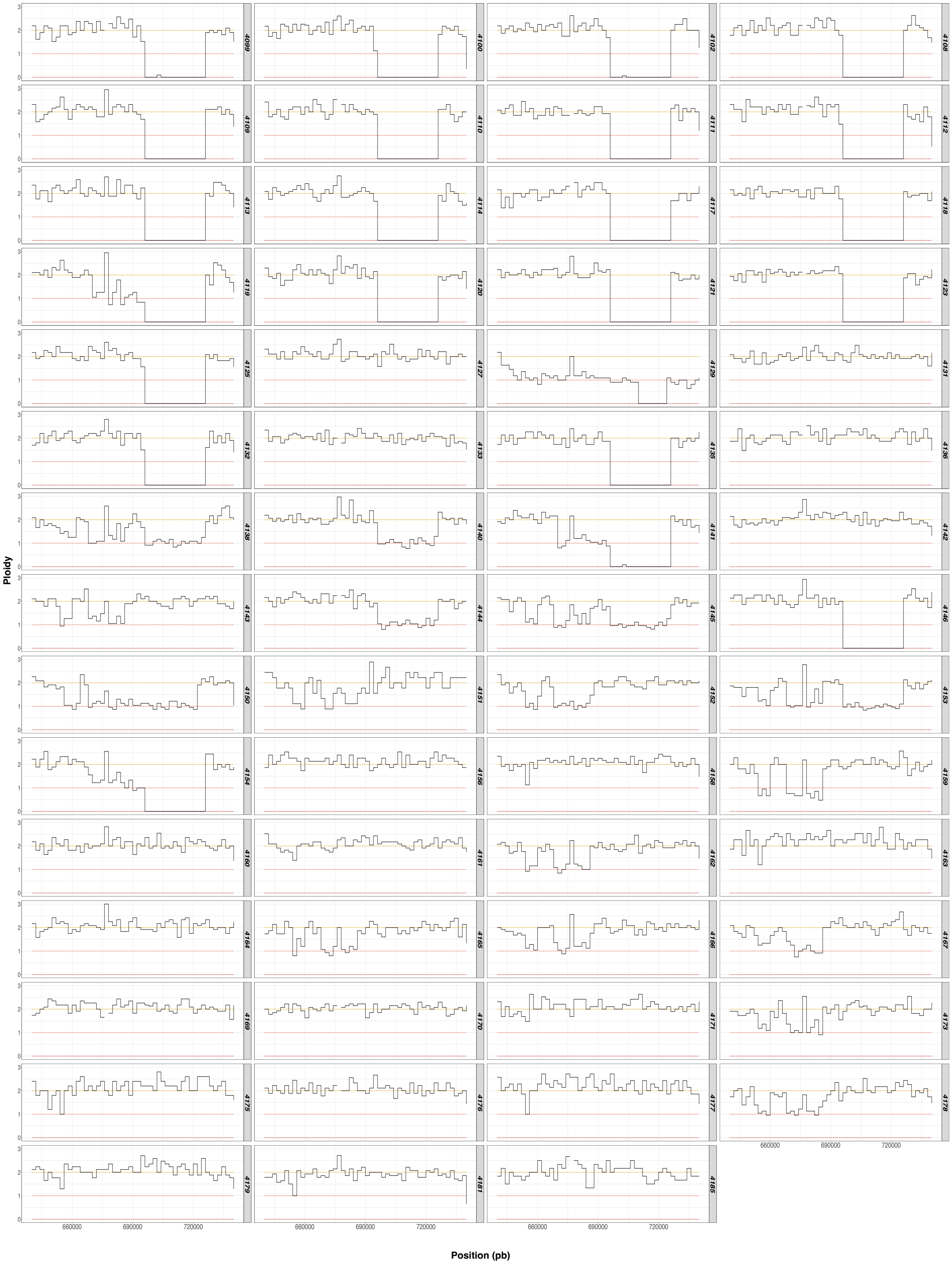

**Fig. S9.** Patterns of coverage in the locus (2/2). The analysis of the locus sequencing coverage between 640 Kpb and 740 Kpb unveils the patterns of deletions impacting the genes g164 and g165. A relative level of coverage of two indicates the presence of sequences both haplotypes, a level of one indicates that the sequence is present in only one of the two haplotypes, 0 copy means a complete deletion of the sequence in both haplotypes.

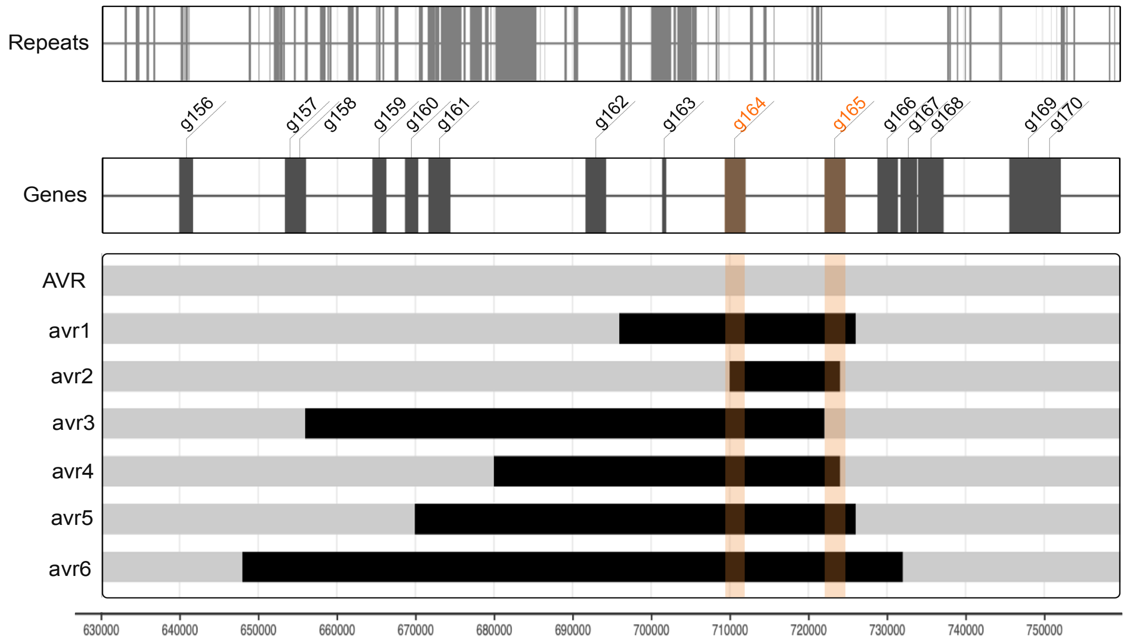

**Fig. S10.** Representation of the seven alleles identified in the 123 *P. viticola* trains analyzed in this study. Genes annotations and repeats elements in the deletion area are represented above the alleles representation. The two genes of interest (g164 and g165 are highlighted in orange as well as their relative position in the deletion). The allele avr1 which exhibited a 30 kbp deletion, from 696 to 726 kbp, was the one observed in the reference genome INRA-Pv221. Then, the deletion in avr2 is from 710 to 724 kbp (14 kbp long), from 656 to 722 kbp (66 kbp long) in avr3, from 680 to 724 kbp (44 kbp long) in avr4, from 670 to 726 kbp (56 kbp long) in avr5 and finally from 648 to 744 kbp (96 kbp long) for avr6.

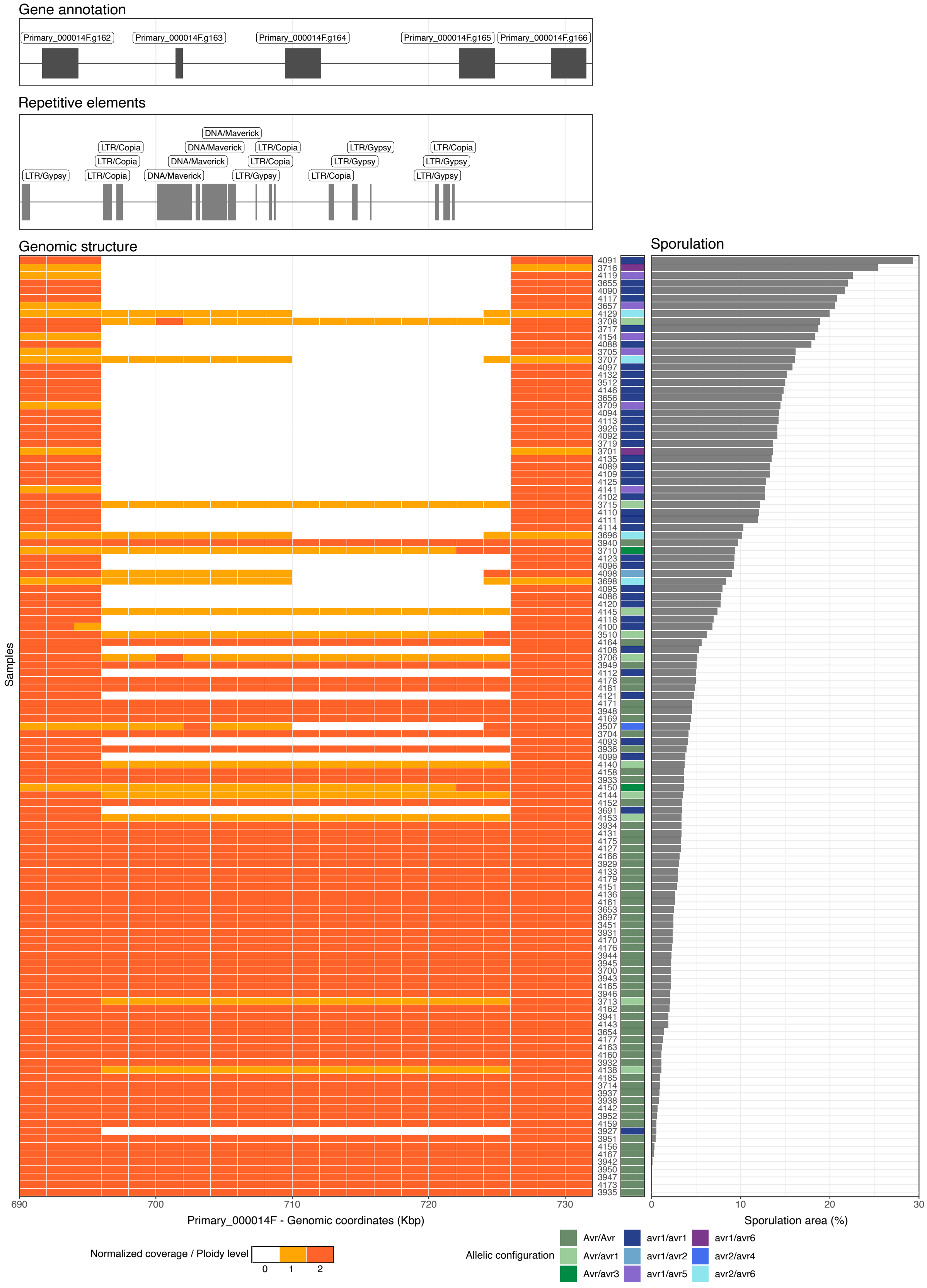

**Fig. S11.** Genomic structure at the avrRpv3.1 locus for each *P. viticola* strains associated with their phenotype (sporulation area on Regent). Gene and repetitive elements annotation are represented above the figure. Genomic structure is represented in the heatmap matrix. The individuals are represented in rows and bins of 2 Kbp in colomn. The ploidy level per bins is represented by the color in each cells: in orange the region is diploid, in yellow it is haploid meaning the deletion happens in only one of the haplotype, in white the deletion happens in both haplotypes. For each rows, at the right of the matrix, a color code indicates the allelic configuration. The sporulation area per strain on Regent in represented on the barplot.

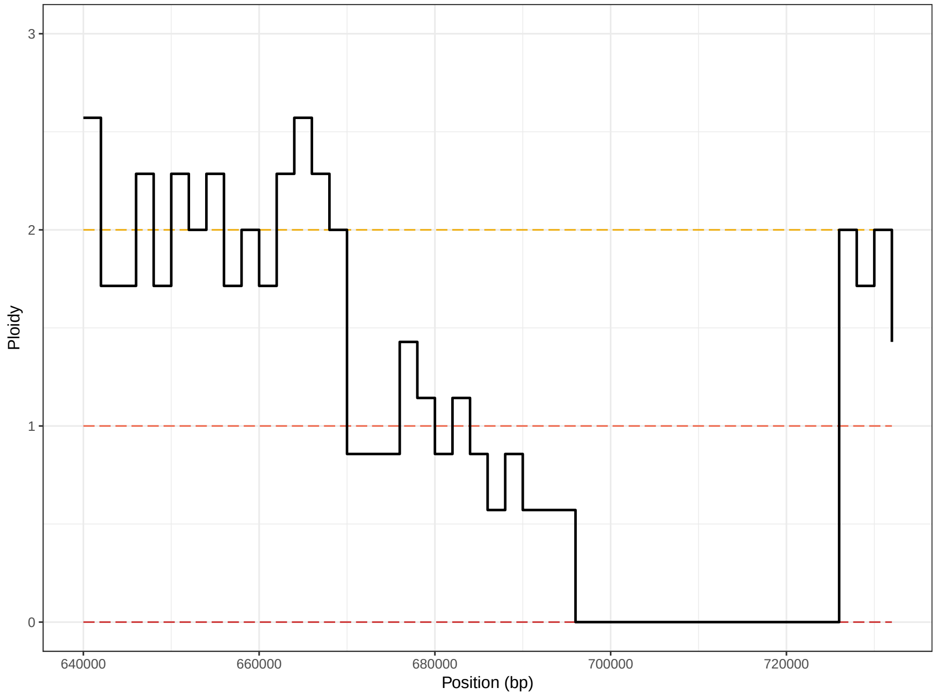

**Fig. S12.** Patterns of coverage in the locus avrRpv3.1 of the strain "avrRpv12-/3-" from (21). The analysis of the locus sequencing coverage between 640 Kpb and 740 Kpb unveils the patterns of deletions impacting the genes g164 and g165. A relative level of coverage of two indicates the presence of sequences both haplotypes, a level of one indicates that the sequence is present in only one of the two haplotypes, 0 copy means a complete deletion of the sequence in both haplotypes.

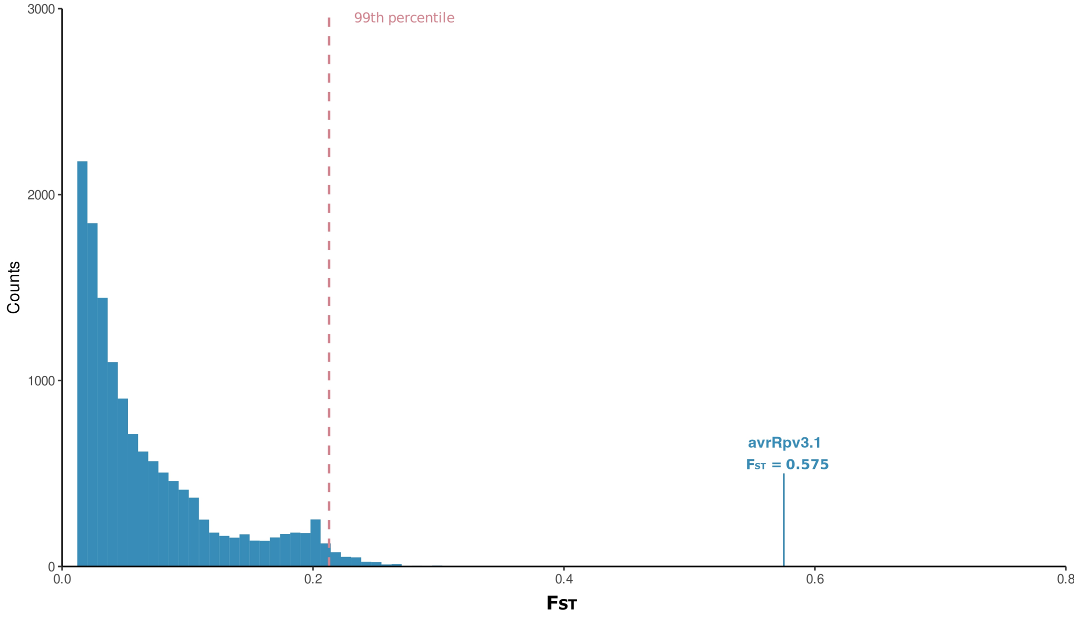

**Fig. S13.** Distribution of Weir and Cockerham FST across the entire contig and the specific FST value computed for the avrRpv3.1 locus indicated by a blue line. The threshold value of the 99th percentile is represented by the red dotted line.

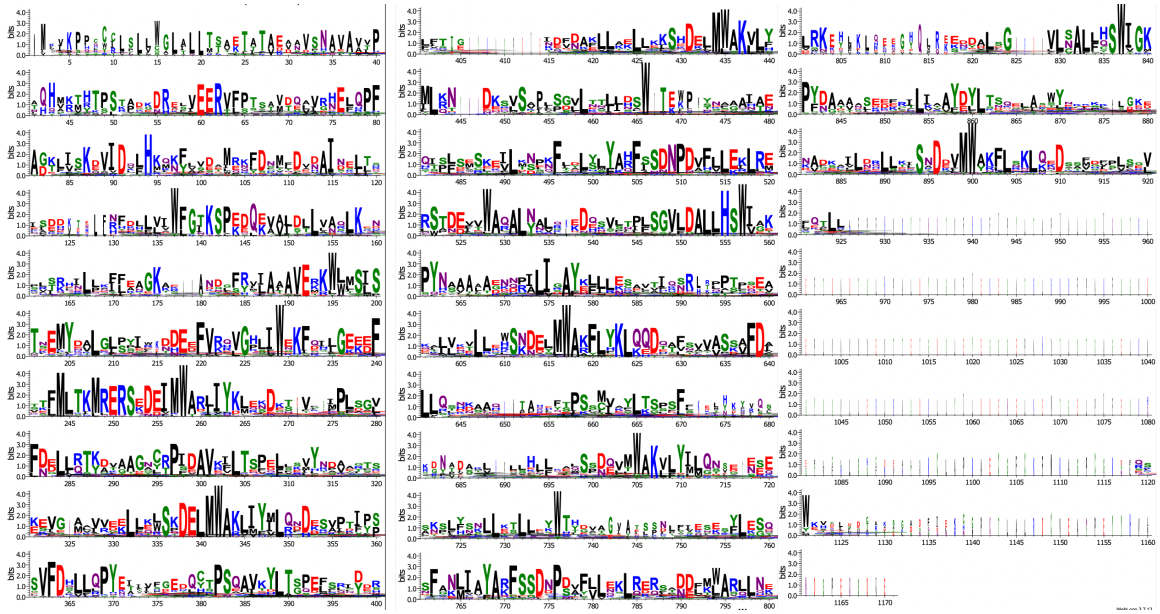

**Fig. S14.** Conserved amino-acid motifs in avrRpv3.1-like proteins

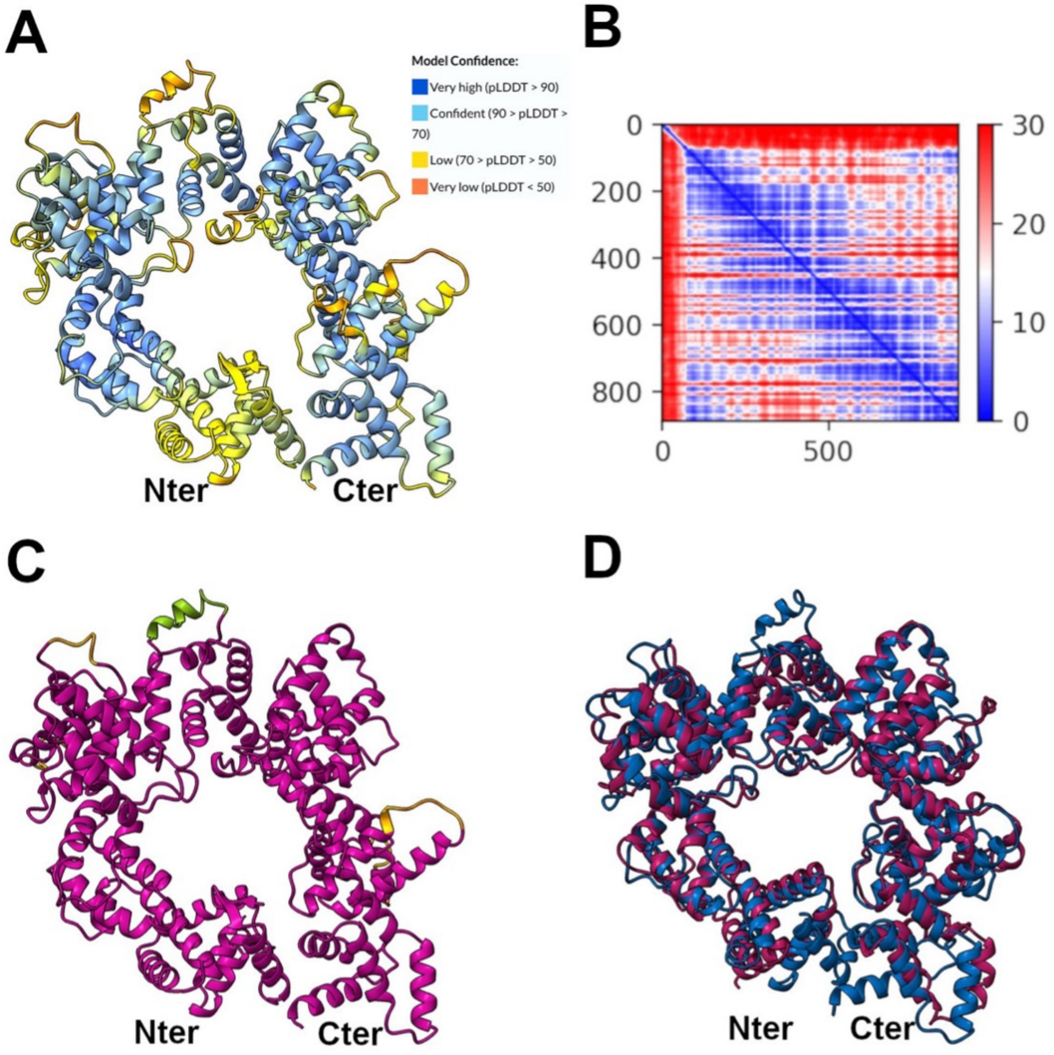

**Fig. S15.** Confidence metrics of the g166 predicted tertiary structure. (A) Predicted structure of g166 colored by per-residue predicted local distance different test (pLDDT) score. The first 75 amino acids have been omitted because of very low confidence. Meaning of coloring is indicated as provided by AlphaFold (https://alphafold.ebi.ac.uk). Values of pLDDT above 70 indicate good backbone prediction. (B) Predicted aligned error (pAE) score of the g166 predicted structure. The lower the score the more confident the prediction. (C) Predicted structure of g166 colored by pAE. Different colors indicate sets of residues with relatively low pairwise pAE values. (D) Superimposition of the predicted tertiary structures of g166 (blue) and a protein from P. halstedii showing 34% identity and 51% similarity to g166 (XP_024580266.1, purple). The first 85 amino acids from both proteins, as well as amino acids from 811 to 915 of the P. halstedii protein, have been removed for clarity. The structure of the P. halstedii protein was retrieved from the AlphaFold Protein Structure Database (https://alphafold.ebi.ac.uk/entry/A0A0P1AQR5).

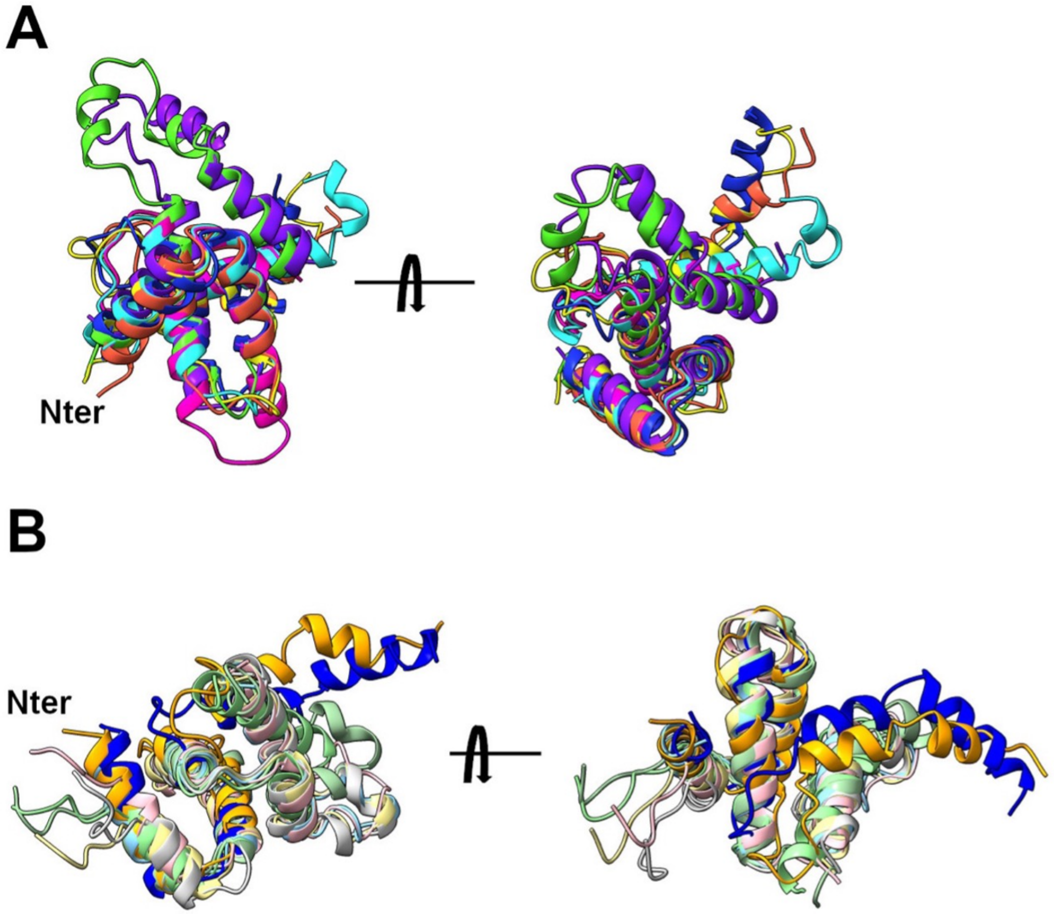

**Fig. S16.** g166 structural domains do not fit the LWY domain fold. The predicted tertiary structure of g166 was divided in structural modules based on the PSR2 LWY domain structure, starting at the N-terminus with the alpha-helices corresponding to the HB1 of the LWY domain. (A) Superimposition of g166 LWY modules 2 to 8. Note overlap at the N-terminus and lack of overlap at the C-terminus. Coloring (2 to 7): orange, yellow, green, cyan, blue, purple, magenta. (B) Superimposition of PSR2 LWYs 2 to 7 and g166 LWYs 3 (yellow) and 6 (blue). Only two g166 LWYs are presented for the sake of clarity. Coloring of PSR2 LWYs (2 to 7): light sky blue, khaki, sea green, silver, light pink, light green.

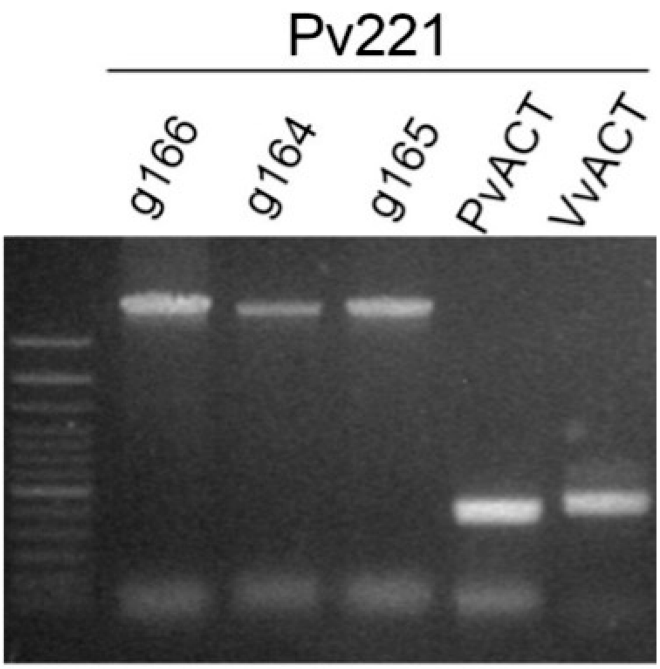

**Fig. S17.** PCRs on genomic DNA from *P. viticola* strain INRA-Pv221. PCRs were performed with primers spanning the complete sequence of each gene, excepting the sequence coding for the signal peptide. PCRs with primers amplifying fragments of the actins from *P. viticola* (PvACT) and *V. vinifera* (VvACT) were used as controls. The identity of the fragments from INRA-Pv221 was confirmed by sequencing. Amplicon sizes: g164-g165-g166: 2600 bp, PvACT 480 bp, VvACT 550 bp. Ladder: Thermo Fisher 100 bp ladder.

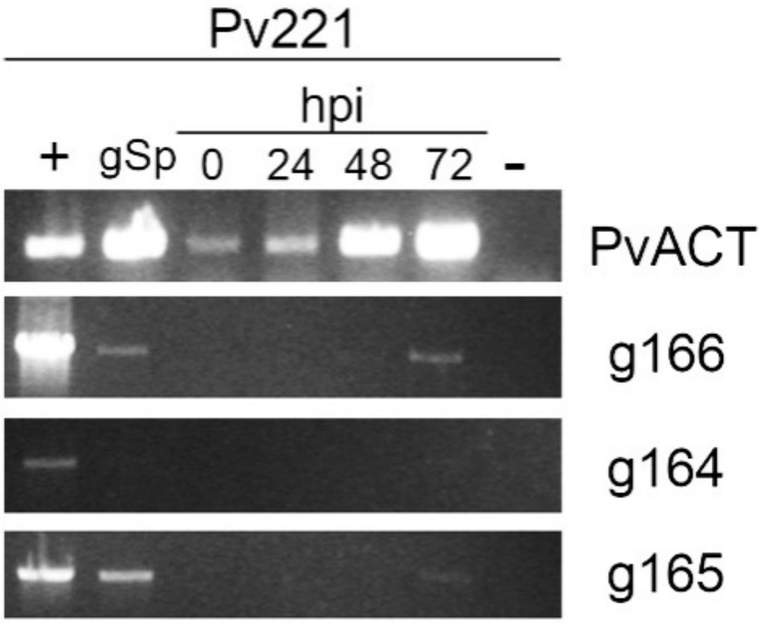

**Fig. S18.** Semi-quantitative RT-PCR showing expression of g164, g165 and g166 in germinated spores (Sg) and infected tissues at 0-, 24-, 48- and 72-hours post-inoculation (hpi), for *P. viticola* strain INRA-Pv221. *P. viticola* Actin (PvACT) expression reveals pathogen biomass and illustrates progression of infection. (+): Genomic DNA, PCR positive control. (-): H2O, PCR negative control. Amplicon sizes: g164-g165-g166: 2600 bp, PvACT 480 bp.

**Table S1.** Statistics of the INRA-Pv221 genome reassembly

| | **Primary** | **Haplotigs** |
|---|---|---|
| **Cumulative length (bp)** | 80,582,756 | 53,611,042 |
| **Number of sequences** | 252 | 745 |
| **GC percentage** | 45.8% | 45.6% |
| **Repetitive content percentage** | 13.4% | 12.5% |
| **Gene loci** | 23,602 | 15,642 |
| **Average sequence length (bp)** | 319,773 | 71,961 |
| **Median sequence length (bp)** | 106,188 | 39,055 |
| **Minimum sequence length (bp)** | 12,273 | 1,266 |
| **Maximum sequence length (bp)** | 3,169,009 | 1,017,603 |
| **N50 Length (bp)** | 825,835 | 134,403 |
| **N50 Index** | 29 | 120 |
| **N90 Length (bp)** | 115,279 | 29,507 |
| **N90 Index** | 116 | 458 |
| **Complete BUSCOs*** | 93.6% | 70.9% |

*BUSCO v2.0, dataset alveolata_stramenophiles

#### supplementary data set

Please note that the supplementary data sets are not available in the current version and will be made available upon publication.

- SI Dataset 1: The origin information of the 136 *P. viticola* strains used in this study and, when available, their phenotype and genotype.
- SI Dataset 2: Multi-alignment of the 11 avrRpvr3.1-like genes and 3 *P. halstedii* effector genes.
- SI Dataset 3: Fasta file of the 11 avrRpvr3.1-like genes and 3 *P. halstedii* effector genes.
- SI Dataset 4: Primers used for PCR and the transient essay.
- SI Dataset 5: RNAseq analysis results
